# Areas activated during naturalistic reading comprehension overlap topological visual, auditory, and somatotomotor maps

**DOI:** 10.1101/037176

**Authors:** Mariam R. Sood, Martin I. Sereno

**Affiliations:** Department of Psychological Sciences, Birkbeck, University of London, Malet Street, London WC1E 7HX; Experimental Psychology, Division of Psychology and Language Sciences, 26 Bedford Way, London WC1H 0AP

## Abstract

Cortical mapping techniques using fMRI have been instrumental in identifying the boundaries of topological (neighbor-preserving) maps in early sensory areas. The presence of topological maps beyond early sensory areas raises the possibility that they might play a significant role in other cognitive systems, and that topological mapping might help to delineate areas involved in higher cognitive processes. In this study, we combine surface based visual, auditory, and somatomotor mapping methods with a naturalistic reading comprehension task in the same group of subjects to provide a qualitative and quantitative assessment of the cortical overlap between sensory-motor maps in all major sensory modalities, and reading processing regions. Our results suggest that cortical activation during naturalistic reading comprehension overlaps more extensively with topological sensory-motor maps than has been heretofore appreciated. Reading activation in regions adjacent to occipital lobe and inferior parietal lobe completely overlaps visual maps, whereas most of frontal activation for reading in dorsolateral and ventral prefrontal cortex overlaps both visual and auditory maps. Even classical language regions in superior temporal cortex are partially overlapped by topological visual and auditory maps. By contrast, the main overlap with somatomotor maps is restricted to dorsolateral frontal cortex.

---

Topological (neighbor-preserving) remapping is a key principle of organization of sensory and motor areas within the mammalian brain. In primary sensory and motor cortices, these representations initially reflect the spatial layout of the receptors; for instance retinotopic maps in visual cortex topologically encode retinal locations, tonotopic maps in auditory cortex represent positions along the cochlear hair cell line, which correspond to sound frequency, and somatotopic maps in the somatosensory cortex represent locations on the body surface. Disrupting these maps has been shown to affect subsequent sensory processing and behaviour (Sperry, 1944; Kaas, 1997). Traditionally it has been assumed that topological mapping was limited to lower level cortex (e.g., Hebb, 1949). But the entire cortex is characterized by the overwhelming predominance of local connections (Lund et al., 1993; Schmahmann and Pandya, 2009; Ercsey-Ravasz et al., 2013). The last few decades of electrophysiological investigations in monkeys and neuroimaging research on humans has shown that topological organization extends well into higher-level cortical areas (Felleman and Van Essen, 1991; Sereno and Allman, 1991; Wandell et al., 2007; Huang and Sereno, 2013). In contrast to lower-level maps, however, localized activity in higher-level maps is affected as much by spatial attention as by the spatial characteristics of the stimuli (Saygin and Sereno, 2008). In frontal cortex, there is evidence that topological maps may serve as a convenient method of allocating working memory, or maintaining pointers to specific content (Hagler and Sereno, 2006), even for tasks not overtly referencing position; for example, the exact areas that showed *more* robust activity during an *identity* two-back task (in which location was ignored), than during a *location* two-back task (in which identity was ignored) turned out to contain retinotopic maps. A significant role for topological maps in other complex mental operations has been suggested before (e.g., Simmons and Barsalou, 2003; Thivierge and Marcus, 2007), but direct neuroimaging evidence supporting this idea has been scant, and mostly confined to somatosensory cortex.

In the study presented here, we used fMRI to directly assess the extent of overlap between cortical regions involved in reading comprehension and those that have a topological sensory or motor map. It is generally assumed that the activation observed in temporal and frontal areas during reading falls beyond the bounds of sensory-motor maps, but this assumption has not been explicitly tested across all modalities in the same group of subjects. Recent advances in cortical surface-based mapping techniques have had less exposure in the language literature; for example, ‘sensorimotor’ regions are often defined only by anatomical features or by basic, non-attention demanding tasks. In this study, we look at the full extent of topologically mapped sensory-motor regions that can be reliably detected using retinotopic, tonotopic and somatomotor mapping and assess where and by how much they intersect with brain regions involved in reading comprehension in the same subjects. To this end, each subject in our study participated in four separate fMRI sessions, where the sessions comprised a naturalistic reading comprehension task, retinotopic mapping, tonotopic mapping, and somatomotor mapping. On the methodological front, we employed a fully surface-based group analysis as opposed to volume-based group analyses commonly used in language studies (merely displaying a 3-D averaged result on an average surface gains none of the benefits of surface-based averaging). The cerebral cortex has the topology of a 2-D sheet. Many relevant dimensions (e.g., retinotopy, somatotopy, tonotopy) vary much more rapidly tangential to the cortical surface than they do perpendicular to the cortical surface, through the several millimeters of cortical thickness. Distances measured in 3-D space between two points – but also used in standard pre-fitting 3-D smoothing – can substantially underestimate the true distance along the cortical sheet due to its folded nature (Fischl et al., 1999a,b). This artifactual within-subject blurring is then made worse by 3-D averaging of between-subject variability in the secondary crinkling of cortical folding patterns. Surface-based techniques make it possible to restrict smoothing to directions parallel to the cortical sheet, and to employ inter-subject 2-D alignment based on the patterns of sulci and gyri after secondary crinkles have been removed, which reduces both kinds of artifactual blurring and improves cross subject averaging (Fischl et al., 1999b); this also provides a less biased estimate of overlap. Finally, siting language regions with respect to topological cortical maps provides a more precise way to compare activations across individuals and groups as well as studies. This is particularly important for refining functional localization in less well-understood regions such as frontal cortex.

## Materials and Methods

### Subjects

20 right-handed native English speakers (9 women) participated in this study. The mean age was 28 (ranging from 19 to 58). All participants were neurologically healthy with normal or corrected to normal vision and normal hearing capacity. The experimental protocols were approved by local ethics committees and participants gave their informed written consent prior to the scanning session. The study required each participant to take part in four separate fMRI experimental sessions: Reading task, Retinotopic mapping, Auditory mapping and Somatomotor mapping. All 20 participants took part in Reading task and Retinotopic mapping experiments. 18 of the same participants took part in Auditory mapping and 17 of the same participants took part in Somatomotor mapping.

### Experimental Stimuli and design

#### Reading Experiment

The reading experiment consisted of a naturalistic reading task where comprehension blocks contained a short narrative passage in English. The experiment was a random-order block design with 3 conditions and a central fixation screen. Each condition block was 16 sec long. During the experimental condition, a passage in English was presented one word at a time for 16 sec (64 words in total, average rate 4 words/sec). Each word was briefly presented on the screen at its natural reading position within the text. All other words appeared as grayed rectangles. The space between successive words/rectangles was set to 35% of the average word length. The exact display time of a particular word was made a piecewise linear function of the rendered word length. Vera.ttf, a san-serif typeface was parsed by FreeType 2.4.11 and rendered and measured in OpenGL using FTGL 2.1.3. The minimum display duration of a word was clamped to 175 msec, which was 70% of the average word duration of 250 msec. Words with widths more than 70% of the average word width were then given linearly increasing display durations with a slope chosen so the total paragraph duration equaled the desired block length (64 sec). This resulted in virtually identical duration/word-length slopes across different 64-word paragraphs. The guided reading experience felt much more subjectively natural when duration was controlled by word length than when a fixed word duration was used. In order to control for low level visual processing, there were two other conditions.

The condition 2 consisted of the 'Hindi' version of the English passage (simple substitution of characters from the font, lekhani_dynamic.ttf, not a translation, so rendered word lengths remained identical) using presentation mode and fixation durations identical to the English condition. In the third condition, Dot, a 0.5 deg visual angle dot instead of a word briefly replaced each of the grayed rectangles, again with the same varying fixation durations and rectangle lengths. The baseline condition (condition 0) presented a single central fixation dot. The experiment consisted of 4 runs, where each run was comprised of 32 blocks presented in a random order. Participants, who had no familiarity with ‘Hindi’ were instructed to do their best to comprehend the English passages in all 4 runs and to follow the Hindi script 'words', or dot in the other two conditions. The passages were self-contained and unrelated to each other. The level of comprehension achieved for the English passages was measured with a questionnaire afterward. Participants were informed of this quality control process before the scan. The stimulus presentation technique used here, where each English word, Hindi word or dot was briefly presented in its natural reading position with all other words grayed out served several purposes. In the spirit of the classic attention study by Posner (1980), the subject’s exogenous attention is automatically drawn towards each newly highlighted position, where the word or dot appears, in a manner very similar to natural reading. The subjects moved their eyes along with the highlighted word and reported a naturalistic reading experience, which was aided by a naturalistic (word-length dependent) fixation duration. Additionally this presentation mode also ensured that participants made controlled eye saccades (and extremely similar eye movements across conditions) as opposed to uncontrolled eye movements if the passages were presented in its entirety. In order to further ensure that participants stayed attentive and made similar eye-movements for all conditions, they were instructed to press a button when the color of an English word, a Hindi pseudo word, a dot, or the central fixation dot changed from black to off-white (for an average of 0.25 seconds). The responses to button press events were logged and analyzed to assess the quality of task execution. Button press events were modeled as an extra regressor and used as an additional quality control check during data analysis. The English screen of the experiment and a sample English passage are shown in Figure 1. The stimulus was programmed in C/OpenGL/X11 (stimulus program available on request). An optimized random order of the conditions within each run was generated using AFNI’s (Cox, 2012) RSFgen program.

**Figure 1:**
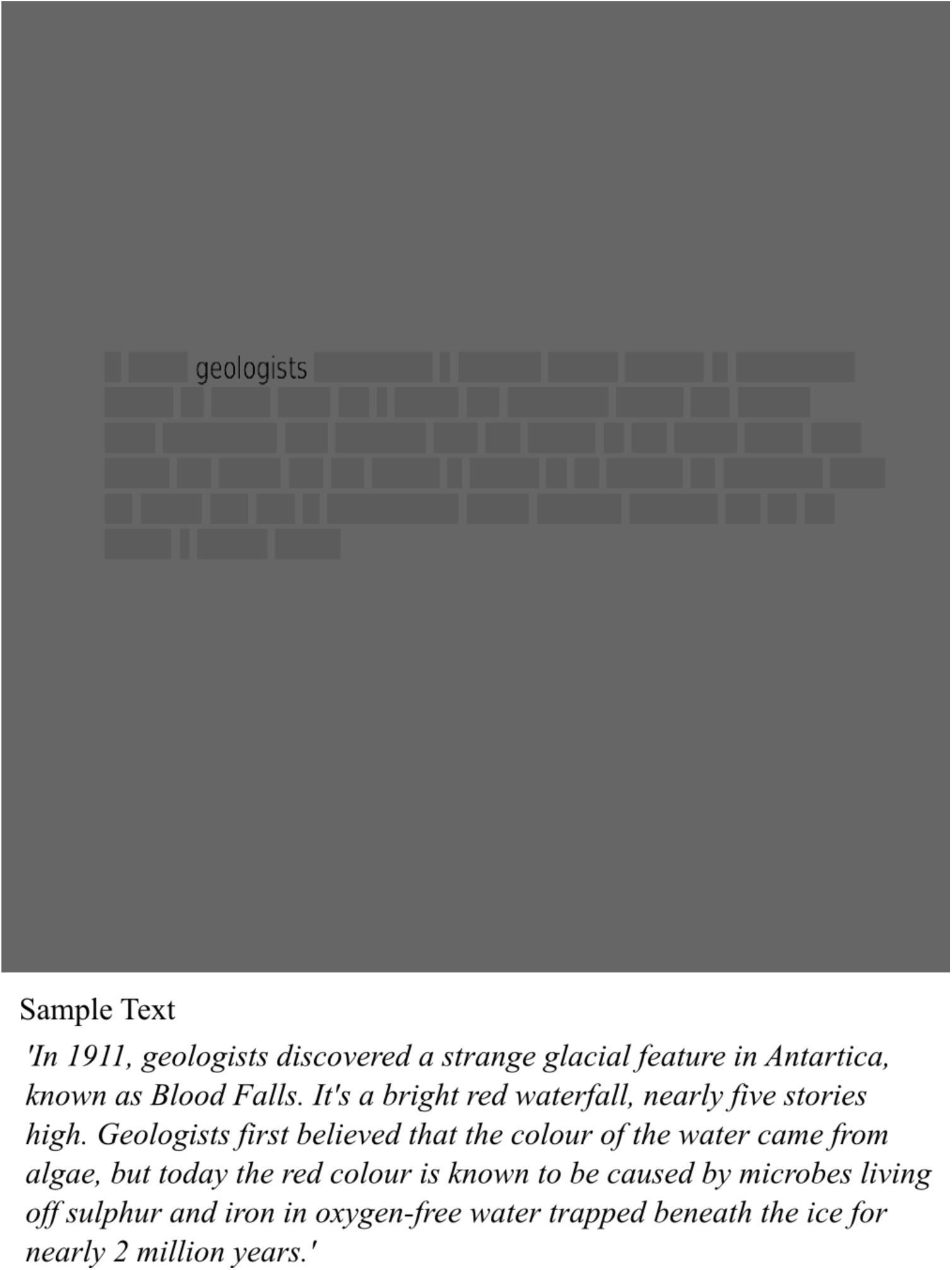
Reading stimulus-English screen and sample English text.

#### Retinotopic Mapping

In the retinotopic mapping experiment, we mapped polar angle using a phase-encoded stimulus very similar to that used in previous recent work by Sereno et al. (2013). The stimulus (Figure 2) consisted of a continuously rotating thin wedge (18 deg wide) populated with a random-colored checkerboard with 35% luminance contrast. The checkerboard was overlaid with white dot fields moving in 500 msec periods of coherent motion that extended slightly beyond the checkerboard wedge to 21.6 deg wide (each new flow period had randomly chosen contraction/dilation and clockwise/anti-clockwise components, dots had 50% average luminance contrast), as well as two simultaneous asynchronous streams of random objects (tiffs of single objects with a transparent background, 0.5 sec duration, 0.1 sec gap) and random black letters (0.4 sec duration, 0.1 sec gap) placed at random eccentricities; both kinds of stimuli were scaled with eccentricity to fit within the confines of the 21.6 deg wide wedge, and their centers were rotated together with the wedge. The objects had an additional random radial (inward or outward, eccentricity-scaled) motion. This polar angle mapping stimulus was designed to evoke activation in the maximum number of lower-level and higher-level visual areas. The participant was presented with the periodic stimulus (64 sec per full rotation cycle, 8 cycles/run). 512 second runs (4 in total) alternated between clockwise and anti-clockwise rotation of wedges. Participants were required to fixate on the center dot at all times. Additionally they were instructed to monitor for occasional numbers (among the letters) and occasional upside down objects (among the right-side-up objects) to maintain a high and continuous level of peripheral attention to the entire wedge during central fixation.

**Figure 2:**
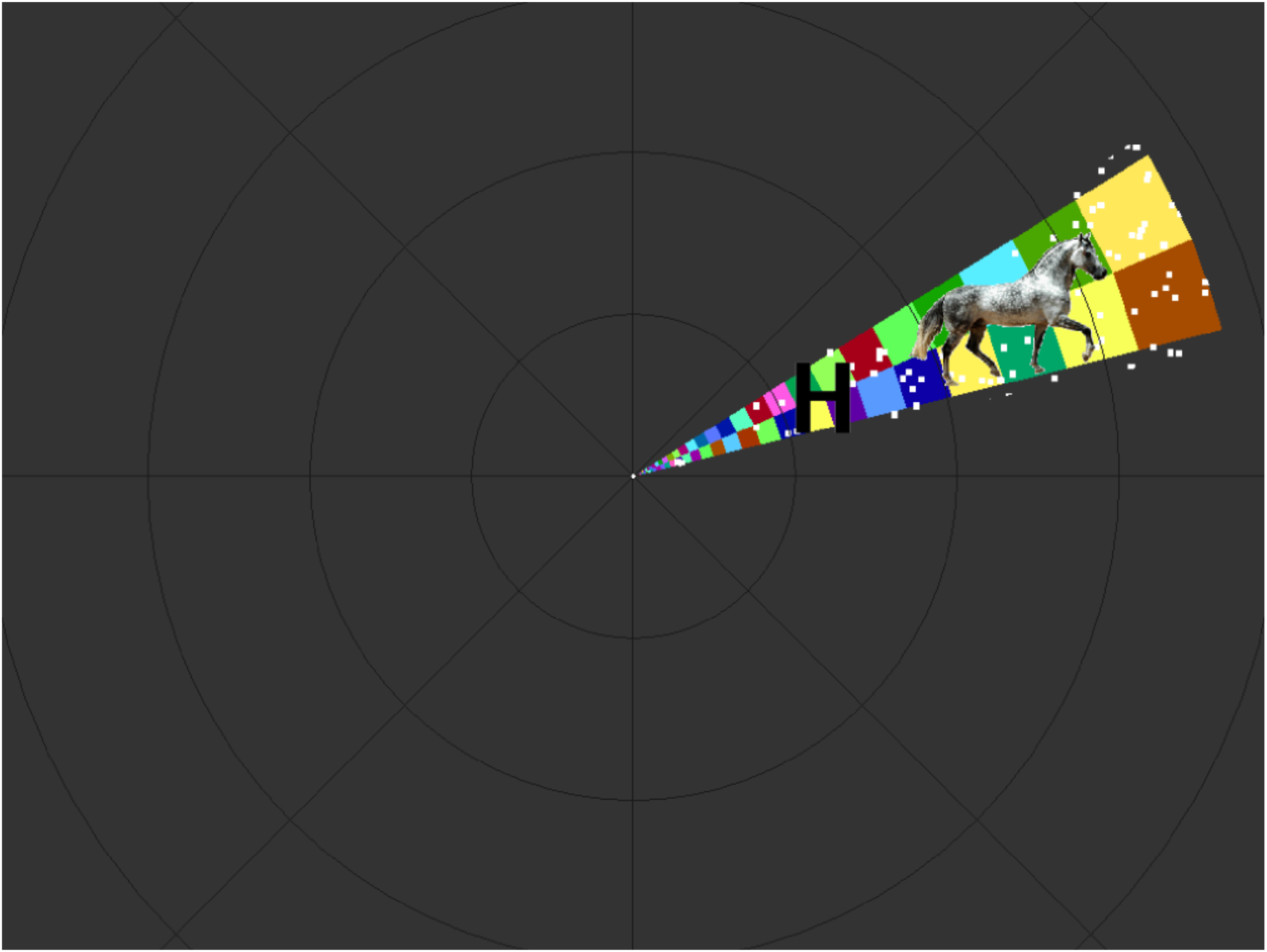
Retinotopic mapping stimulus.

#### Auditory Mapping

In the auditory mapping experiment, we mapped tonotopically organized cortical areas using a frequency-modulated stimulus taken unchanged from the previous study by Dick et al. (2012). The stimulus consisted of bandpass-filtered nonlinguistic vocalizations adapted from the Montreal Affective Voices (Belin et al., 2008), a series of recordings by actors (only male voices were utilized in the stimulus) producing sounds associated with a set of eight emotions. Each run consisted of eight 64 sec band-pass-filtered cycles, where the center filter frequency repeatedly logarithmically ascended from 150 to 9600 Hz, or repeatedly descended from 9600 to 150 Hz (no gap at the wraparound point). The session consisted of 4 runs alternating between ascending and descending frequency sweeps. During scanning, subjects were asked to monitor the stimuli and press a button whenever they heard laughter (laughter targets were distributed non-periodically through the stimulus train). The band pass filter made this task challenging.

#### Somatomotor Mapping

In order to reveal sensory and motor maps in primary and secondary somatosensory and motor cortical areas, participants were asked to move different body parts in response to periodic auditory cues (spoken name of the body part to be moved, rendered using Mac OS X text-to-speech with "Alex" voice) in as gentle and localized fashion as possible. Each run consisted of 8 cycles of 64 sec, and in each cycle participants successively moved 11 different body parts progressing from tongue to toe. Runs (4 in total) alternated between movement cycles in each direction (from tongue to toe and toe to tongue). The stimuli were presented by an in-house C/OpenGL program and the conceptual design is similar to that described in Zeharia et al. (2012). The participants were allowed to familiarize themselves with the auditory cues and practice the controlled movements prior to the scan.

Auditory instructions were: "say tchuh-tchuh"; "say pup-pup"; "crinkle eyebrows"; "touch thumb index"; "wave wrist"; "contract biceps"; "pull in stomach"; "squeeze buttocks"; "contract quads"; "wave ankle"; "rub big toe".

### Experimental set-up

For visual experiments (Reading task and Retinotopic mapping), the stimuli were projected into the bore using an Eiki LC-XG300 XGA video projector onto a translucent direct-view screen at the participant’s upper chest level. All polar angles of the visual field were stimulated out to an eccentricity of at least 54 degrees of visual angle (much larger than the usual 8-12 degrees of all-polar-angles eccentricity achieved when a standard screen is viewed via a mirror). This avoids artifactual periodic modulation of voxels representing visual field locations beyond the outer edge of the stimulus due to surround inhibition. A black matte shroud situated just outside the bore blocked the beam from making low-angle reflections off the top of the bore. The rear of the head coil was elevated with a wooden wedge and thinner bed cushions were used to help naturally tilt the head forward. For auditory and somato-motor mapping, the stimuli were delivered binaurally using in-house safety-enhanced Sensimetrics (Malden) S14 earbuds and cushions. During all scanning sessions, memory foam cushions (NoMoCo Inc.) were packed around the head to provide additional passive scanner acoustical noise attenuation and to stabilize head position. Responses were made via an optical-to-USB response box (LUMItouch, Photon Control, Burnaby, Canada) situated under their right hand.

### Imaging Parameters

Functional images were acquired on a 1.5 T whole-body TIM Avanto System (Siemens Healthcare), at the Birkbeck / University College London Centre for NeuroImaging (BUCNI), with RF body transmit and a 32-channel receive head coil. For the first 16 fMRI sessions, images were acquired using the standard product EPI pulse sequence (24 slices, 3.2x3.2x3.8mm, 64x64, flip=90°, TE=39ms, TR=2sec), while the remaining 64 sessions used multiband EPI (40 slices, 3.2x3.2x3.2mm, flip=75°, TE=54.8ms, TR=1sec, accel=4) (Moeller et al., 2011). Individual scans had 260 volumes for standard EPI and 520 volumes for multiband EPI. To allow longitudinal relaxation to reach equilibrium, 4/8 initial volumes were discarded from each run for standard/multiband EPI. For each imaging session, a short (3 min) T1-weighted 3D MPRAGE ‘alignment scan’ (88 partitions, voxel resolution 1x1x2mm, flip angle=7^o^, TI=1000ms, TE=4ms, TR=8.2ms, mSENSE acceleration=2x, slab-selective excitation) was acquired with the same orientation and slice block center as the functional data, for initial alignment with the high-resolution scans used to reconstruct the subject’s cortical surface. Two high resolution T1-weighted MPRAGE scans (176 partitions, 1x1x1mm, flip angle=7^o^, TI=1000ms, TE=3.57ms, TR=8.4ms) were acquired along with the fMRI sessions for cortical surface reconstruction (using FreeSurfer 5).

### Data Analysis

#### Anatomical image processing

For each subject, the cortical surface was reconstructed with FreeSurfer (version 5; Dale et al., 1999) from the aligned average of the two high-resolution T1-weighted MPRAGE scans. Both mapping data and reading data employ a complex-valued cross-subject surface-based analysis stream that begins by sampling responses and statistics to individual reconstructed cortices (cross-subject 3D averaging was not used at any point in the pipeline).

#### Analysis of phase-encoded mapping data

The first level single subject data from each run was motion corrected and registered to the last functional scan (across all runs) using AFNI’s 3dvolreg (Cox, 2012) program (using heptic interpolation). The functional images were registered with the cortical surface using a multi-stage registration pipeline. This involved first registering the ‘alignment’ scan to the high resolution MPRAGE, and then transferring that registration to the functional EPI images. These steps were done using FSL’s FLIRT tool (version 5; Jenkinson and Smith, 2001; Jenkinson et al., 2002) and were then fine tuned using FreeSurfer’s bbregister (Greve and Fischl, 2010). The final combined and fine-tuned 4x4 registration matrix aligns functional EPI to high-resolution MPRAGE (the matrix actually transforms 3D surface vertex coordinates to 3D EPI block index coordinates). The final registrations (EPI -> hi-res surface MPRAGE) were visually checked to ensure accuracy. The time courses from all the runs were averaged (after time-reversing the runs in the clockwise rotation for retinotopy, the upward frequency sweep for auditory, and the toe-to-tongue movement direction for the somatomotor map). The time reversed scans were time-shifted to compensate for hemodynamic delay before averaging. The averaged time courses were analyzed using linear Fourier methods (Bandettini et al., 1993; Engel et al., 1994, 1997; Sereno et al., 1995; Hagler and Sereno, 2006), which can be exactly recast as a general linear model. Voxels preferentially responding to a particular point in the stimulus cycle will show higher amplitude at the stimulus frequency than at any other 'noise' frequency, after excluding (i.e., linearly regressing out) the 3 lowest temporal frequencies as motion artifact. For retinotopic data, the phase of this vector at the stimulus frequency indicates the polar angle of the stimulus location. For auditory data, the phase indicates a particular point on the stimulus frequency ramp. For somatomotor data, this corresponds to the location of the moved body part. The individual and group analyses utilized a complex-valued cortical surface based stream that was previously described (Sereno et al., 1995; Hagler et al., 2007; Huang et al., 2012) and briefly summarized below.

A fast Fourier transform was first performed on the average time courses of each voxel. An F-statistic value was obtained for each voxel by comparing the power at the stimulus frequency (eight cycles per scan) to the average power at the remaining frequencies after excluding the second and third harmonics of the stimulus frequency and one frequency above and below the first three harmonics. For individual subjects' activations illustrated below, the F-statistic was thresholded at p<0.001 (corresponding to F(2,232)=7.12 for subject 1 who was scanned on standard EPI with 256 time points and F(2,488)=7 for subject 2 and subject 3 who were scanned on multiband EPI with 512 time points). Surface-based cluster size exclusion (Hagler et al., 2007) was used to correct for multiple comparisons with cortex surface clusters smaller than 30 mm^2^ excluded, achieving a corrected p-value of 0.01.

Group analysis of phase-encoded mapping data was performed using the methodology developed by Hagler et al. (2006) in which the real and imaginary components of the signal at the stimulus frequency were averaged across the subjects, preserving any phase information consistent across subjects (this is a vector average, which properly treats wrap around in the circular phase variable). This was performed by projecting each participant’s complex-valued phase-encoded map to the FreeSurfer spherical atlas (using FreeSurfer mri_surf2surf), only performing ten steps of surface-based smoothing (∽3mm FWHM in 2D) before vector averaging across subjects at each vertex in the common surface coordinate system. Second-level surface-based cluster size exclusion was used to correct for multiple comparisons, with vertex level F-statistics thresholded at p<0.01/p<0.05 and cortical surface clusters smaller than 40 mm^2^/92 mm^2^ excluded, achieving a corrected p-value of 0.05. We used the fsaverage "inflated_avg" surface for display (made by averaging inflated surface coordinates) instead of the fsaverage "inflated" surface (made by averaging folded surface coordinates and then inflating the average) because inflated_avg represents original average surface area better. This is because folding variations (sulcal crinkles) are removed before surface-averaging, making inflated_avg, more appropriate for displaying surface-averaged data (mesh defects in the north and south icosahedral poles of FreeSurfer5's inflated_avg were corrected before using it for display. Corrected and flattened inflated_avg surfaces are available here: http://www.cogsci.ucsd.edu/∽sereno/.tmp/dist/csurf/fsaverage-adds.tgz).

#### Analysis of Reading comprehension data

For the reading experiment, single subject fMRI data was motion corrected and skull stripped using FSL tools (MCFLIRT and BET). First level fMRI analysis was carried out by applying the General Linear Model (GLM) within FEAT using FILM prewhitening (FSL, version 5) with motion outliers (detected by fsl_motion_outliers) being added as confound regressors if there was more than 1 mm motion (as identified by MCFLIRT). A small number of scans with excessive motion above the threshold of 1 mm were excluded from analysis. High-pass temporal filtering of the data and the model was set to 100 seconds based on the power spectra of the design matrices (estimated by cutoffcalc; part of FSL). Three main explanatory variables were modeled and controlled: Reading English text, viewing Hindi text and viewing dot ‘text’. Button press responses to target color change were modeled as the fourth regressor. In order to capture slight deviations from the model, temporal derivatives of all explanatory variables convolved with FEAT's double gamma hemodynamic response function (HRF) were included. The registration from functional to anatomical (6 DOF) and standard space (12 DOF) was first done using FSL’s FLIRT and further optimized using boundary based registration (bbregister; FreeSurfer) similar to the procedure for the phase-encoded mapping data. A fixed effects analysis was performed across runs from an individual subject (usually 4 runs unless a run was excluded due to excessive motion/ lack of attention as evidenced by poor performance on qualitative assessment/ lack of response to the targets) to get group FEAT (GFEAT) results of first-level contrast of parameter estimates (COPEs) and their variance estimates (VARCOPEs) in the standard space. Across-subject group analysis was then carried out on the cortical surface using FreeSurfer tools. The GFEAT results of each subject were first sampled to individual cortical surfaces and then resampled to the spherical common average reconstructed surface (fsaverage). Surface based spatial smoothing of 3 mm FWHM was applied on the icosahedral sphere. A mixed effects GLM group analysis was performed on the average surface using the mri_glmfit program from FreeSurfer. Significance maps (p<0.01) were then corrected for multiple comparisons with cluster-based Monte-Carlo simulations with 10,000 permutations of white Gaussian noise using the FreeSurfer program mri_glmfit-sim. Finally corrected significance values (p<0.05) of reading activation were displayed on the average surface.

The single subject raw data was not spatially smoothed (both for phase-encoded analysis and reading analysis). As with the phase-encoded data, a 10 step (∽3mm FWHM) surface level smoothing was applied for final illustration of the results. Hence, 3D Gaussian random field based cluster correction provided by FSL was not appropriate for multiple comparison correction of the reading data. We have instead used the surface based cluster correction similar to that employed for phase-encoded data analysis. The GFEAT results were sampled to their respective anatomical surface, thresholded at p<0.001 (Z=3.09) and corrected for multiple comparisons with cortex surface clusters smaller than 30 mm^2^ excluded, achieving a corrected p-value of 0.01.

The target (font color change) presentation timings and the button press events were logged during the experiment and analyzed to assess the performance of the task. A target was considered as detected if there was a response (a key press event) within 1 second after the colour change event had ended. For each participant, the number of targets detected and the mean response time in each condition were calculated. The cross-subject mean response time and target detection rates were assessed for significant differences across conditions.

#### Overlap analysis

All overlaps were calculated using "original vertex-wise area" in FreeSurfer. Original vertex-wise area in FreeSurfer is defined as the sum of 1/3 the area of each adjacent triangular face on the FreeSurfer "white" surface (refined gray/white matter boundary estimate). That single-vertex sum is not exactly constant across vertices because of slight non-uniformities in the final relaxed state of the surface tessellation. However, the sum of vertex-wise areas over a connected region of vertices exactly represents the summed original area of the enclosed triangles (plus the 1/3 fraction of triangles associated with the boundary vertices; along a straight edge of vertices, this last contribution corresponds to half of the area of the triangles just beyond the edge). The minimum areal increment that can be measured is roughly the average original vertex-wise area, which is ∽0.6 sq mm.

## Results

We first discuss the results obtained for each experiment in the study individually, followed by the overlap analysis results of reading activation with topological visual, auditory and somatomotor maps. A separate section is included to discuss several new and previously unreported sensory maps observed during the mapping experiments.

For the result figures corresponding to individual experiment results, we illustrate the amplitude of the vertex wise response for both reading experiment and phase-encoded maps (ignoring phase) for a stairstep of t-values. This is followed by figures illustrating single modality phase hue maps for sensory-motor maps where hue is used to indicate the map coordinates.

The overlap figures use transparent overlays to indicate reading activations over single modality phase hue maps, finally leading up to a summary outline figure containing all four kinds of data. The language contrast used for overlap analysis is English vs. Hindi. Brain activation observed for English vs. Hindi contrast partially overlapped with retinotopic, tonotopic and somatomotor maps. For clarity, in overlap figures, only positive activation after thresholding and cluster correction is shown. The overlap results for different modalities are illustrated for several individual subjects and then for the group as a whole. For the cross-subject average, the sensory-motor maps are illustrated for two separate vertex thresholds; p<0.05 (lower threshold) and p<0.01 (higher threshold), corrected for multiple comparisons using cluster thresholding at p<0.05. Because the phase-encoded analysis effectively spreads the same amount of imaging data over a larger number of different effective conditions (e.g., different polar angles, sound frequencies, body parts) than the Reading experiment does, the regression analysis carried out on reading data will have more power. Hence the Reading data illustrated here uses a vertex threshold of p<0.01 in all cross-average images. For the individual subjects, results are illustrated at a higher vertex threshold of p<0.001, corrected to p<0.01 for both reading and mapping data due to the higher amount of noise present in single subject data.

The individual subject data are illustrated here to show that in general the pattern of activity was similar to cross-subject average. The majority of subjects showed reading activation patterns similar to subject-1 and subject-2, while subject-3 had more restrained reading activation. All three of the individual subjects illustrated here had superlative comprehension task performance (vivid post-scan explanations correctly citing most of the text concepts presented to them during the task); all were in their early 20’s.

Among the 20 subjects who took part in reading experiment, data from 3 subjects were excluded from the analysis owing to unsatisfactory performance in the target detection task and/or assessment of poor comprehension following the session and/or excessive movement (>1 mm). The activation for the target detection regressor (button press regressor) was used as an extra quality check to decide whether the subject performed the task as per the instructions during each run. All our included subjects had comparable performance across conditions in the target detection task and motor activation for the target regressor. Among the 17 (of the same) subjects who took part in somatomotor experiment, data from 2 subjects were excluded owing to high stimulus correlated head motion during the task. No exclusions were made for retinotopic and tonotopic experiments.

### Target detection response

Figures 3 shows the target detection response results based on the performance of the included subjects. The average response time for the group when the target occurred in English, Hindi, Dot and Fixation conditions are depicted in Figure 3A. On average, participants took 0.47 seconds ± 0.01 (SEM) to respond to the target when it occurred in English/Hindi conditions and 0.42 seconds ± 0.01 (SEM) and 0.44 seconds ± 0.02 (SEM) when they occurred in Dot/ Off conditions. A Wilcoxon matched-pairs signed-ranks test indicated no significant differences between ‘English’ (median= 0.45 seconds) and ‘Hindi’ (median=0.46 seconds) (Z=-1.681, p=0.098). The differences between ‘English’ and ‘Dot’ (median=0.42 seconds) and ‘English’ and ‘Off’ (median = 0.40 seconds) conditions were found to be significant (Z=-2.96, p=0.002 for English-Dot and Z=-2.02, p=0.043 for English-Off).

**Figure 3:**
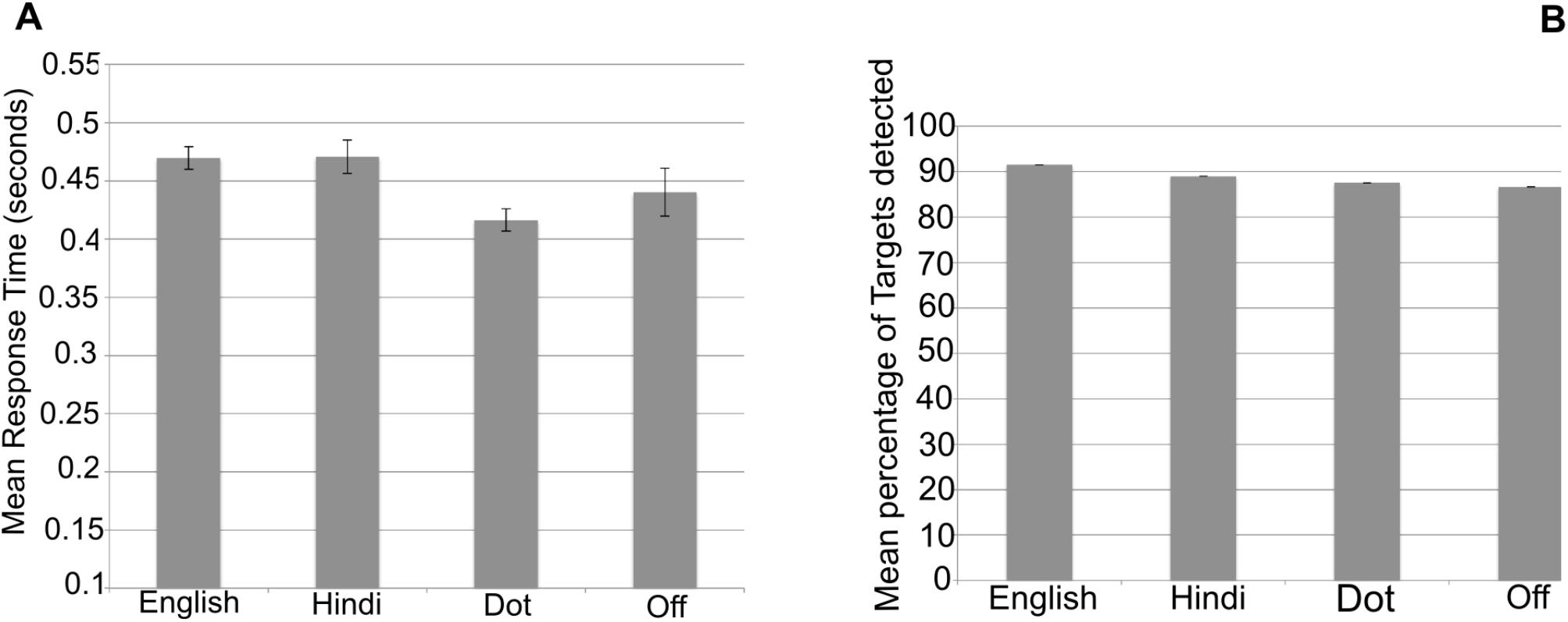
Target detection Performance results (Reading Experiment). Error bars represent one standard error from the mean.

Figure 3B depicts the average target detection success rate across all participants for different conditions. Across all runs, there were 20 targets for the English condition, 18 for the Hindi condition, 23 for the Dot and 7 for Off condition. All participants except one (see below) had comparable detection performance irrespective of the condition in which the target occurred. One participant did not respond to all 7 targets that fell on the fixation, though she consistently performed in all other conditions. All other quality measures were satisfactory for this subject and her data suggested that she did fixate during Off condition (the fovea had no activation in English vs. Fixation). We have therefore conservatively considered her data as valid. On average, the mean success rate for detecting targets when they occurred in English, Hindi, Dot and Off were 91.6%, 89.1%, 87.5% and 86.6% respectively. The average for Off condition is less mainly because of one outlier, the above mentioned subject. Excluding her data, the average success rate for Off condition was 92%. A Wilcoxon matched-pairs signed-ranks test was carried out to assess statistical significance of the average success rate. The median success rate for English, Hindi, Dot and Off were 90%, 89%, 87% and 100% respectively. There were no significant differences between English and any of the other conditions (English-Hindi: Z=-1.733, p=0.087; English-Dot: Z=-1.949, p=0.051; English-Off: Z=-0.94, p=0.94).

### Reading Activation

Figure 4 and the top of Figure 5 illustrate the average cross-subject activation for a stairstep of t-values (all beyond a minimum threshold of p<0.05, uncorrected) for each condition (English, Hindi and Dot) relative to fixation (Fig. 4A-C), and those for two main contrasts, Hindi vs. Dot (Fig 4D), and English vs. Hindi (Fig. 5A). The main contrast used to assess reading comprehension is English vs. Hindi. In the later Figures depicting overlap with sensory-motor maps, transparent bright yellow regions outlined in black depict the regions that showed significantly higher activation when reading English compared to Hindi. In all subjects, activation for English > Hindi was more widespread and pronounced in left hemisphere than in right hemisphere, as expected. The right hemisphere regions activated were a mirror image subset of the left hemisphere counterparts.

**Figure 4:**
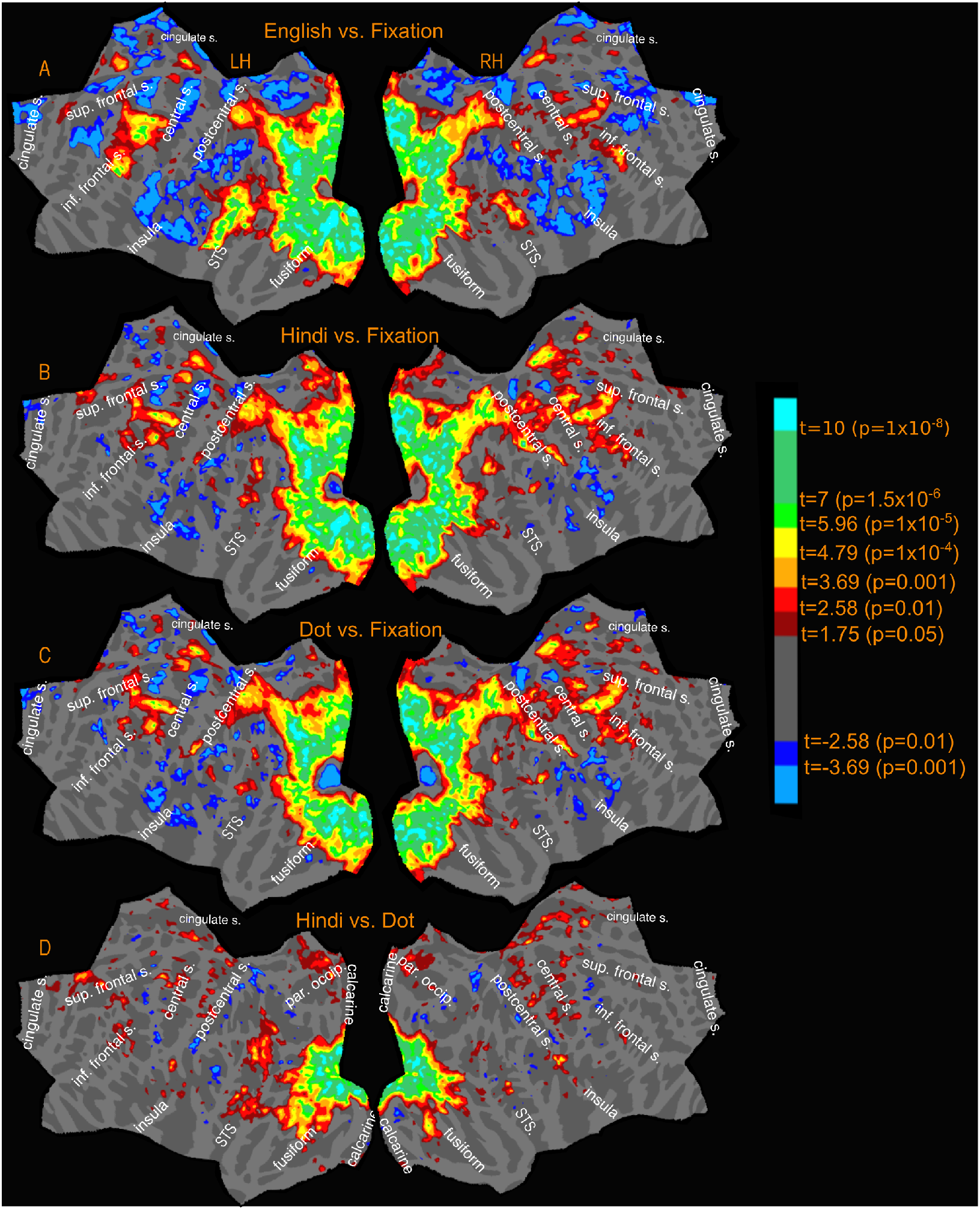
Reading Experiment-Activation amplitude profile (uncorrected) for relevant contrasts.

**Figure 5:**
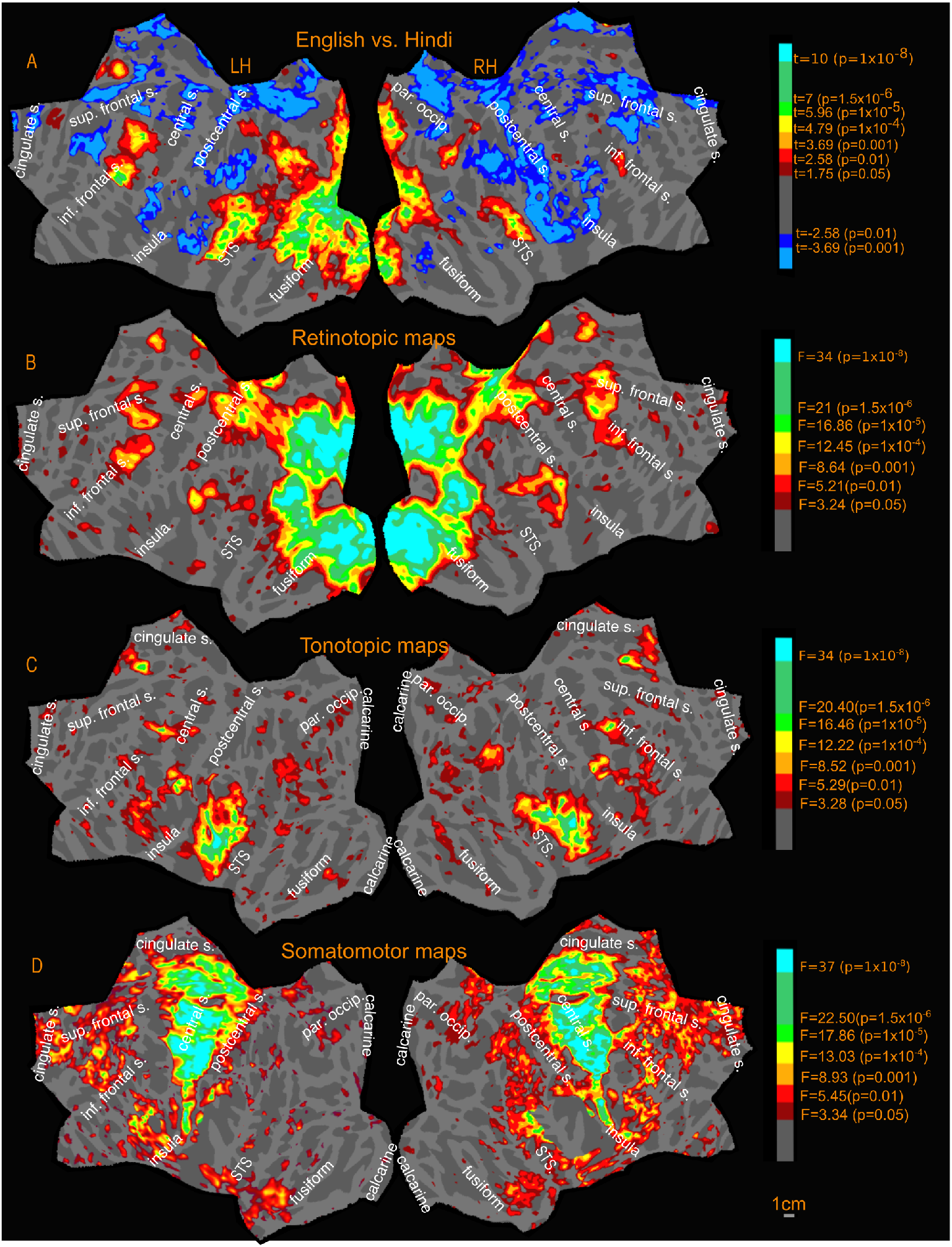
Activation amplitude profile (uncorrected) of Sensory-motor maps and English vs. Hindi contrast.

The cross-subject reading activation for English vs. Hindi (Figure 5A) is spread around 3 main connected regions in the left hemisphere. The first region was anterior to the occipital pole, extending laterally into the occipital cortex with two 'offshoots'. The first offshoot stretched across the intraparietal sulcus, covering regions in the inferior and superior parietal lobules. The second offshoot stretched nearly half the length of the inferior temporal gyrus and extended ventrally onto the fusiform gyrus joining the medial activation that extended from calcarine fissure to the fusiform gyrus. Less extensive but prominent activation was observed in medial regions in the cuneus, precuneus and at the isthmus of the cingulate gyrus. For English, Hindi, and Dot (versus Fixation), roughly the same regions in the occipital and parieral lobe were activated. However, the activation differences were most significant in those regions for English vs. Hindi.

The second main region (English vs. Hindi) is the superior temporal cortex, covering regions along the superior temporal gyrus and sulcus extending well into the middle temporal gyrus and supramarginal gyrus. Except for a small region in posterior STS, the Hindi and Dot conditions do not have any significant temporal lobe activation.

The third region included two distinct frontal regions – one near the precentral sulcus and another near the inferior frontal sulcus in the pars opercularis region. The right hemisphere activation profile is similar but covering a smaller total extent of cortical area. The activation near the precentral sulcus is present for Hindi and Dot conditions as well, with no significant differences in the Hindi vs. Dot contrast (Fig. 4D). The activation observed for English in inferior frontal region near pars opercularis is absent in both Hindi and Dot conditions.

The reading activations of subject-1 and subject-2 were strikingly similar to the cross-subject profile, with activated regions corresponding to the three main regions just described in the left hemisphere. Individual subject-3 (see below) showed the greatest variation from the average, with more extensive activation observed in superior temporal sulcus and middle temporal gyrus, but then less extensive activation elsewhere in the cortex.

### Retinotopic maps

The retinotopic maps (amplitude in Fig 5B, phase in Fig. 6A; see also phase underlays in Figs. 7, 8, and 9) branch out anteriorly from the occipital pole into several 'streams' (numbered 1-5 in black in Fig. 6A). In all phase maps, lower visual field is green, horizontal meridian is blue, and upper visual field is red (all contralateral). One stream extends through area MT into the superior temporal sulcus and reaches the posterior lateral sulcus (leaving a few disconnected regions in between, which join up as threshold is slightly lowered). A second stream stretches along the intraparietal sulcus and arrives at the superior part of the postcentral sulcus. A third stream spreads across the parieto-occipital sulcus (POS) into the medial posterior parietal cortex and precuneus, ending at the cingulate sulcus visual area (Huang and Sereno, 2013). A fourth stream runs across the POS into retrosplenial cortex at the isthmus of cingulate gyrus (iCG) and continues to the edge of cortex just under the splenium of the corpus callosum. Finally, a fifth stream follows the collateral sulcus and fusiform gyrus into the ventral occipitotemporal lobe. A further disconnected set of retinotopic maps, including the frontal eye fields (FEF), frontal poly-sensory zone (Huang et al., 2012), and dorsolateral prefrontal cortex (DLPFC) are found in the frontal cortex. These maps are similar to those reported previously in Huang and Sereno (2013) using a similar stimulus. We also found a previously unreported visual map in the anterior cingulate region, referred to in Figure 6A as anterior cingulate visual area (ACv), which corresponds to human dorsomedial eye-fields (discussed further under New maps).

**Figure 6:**
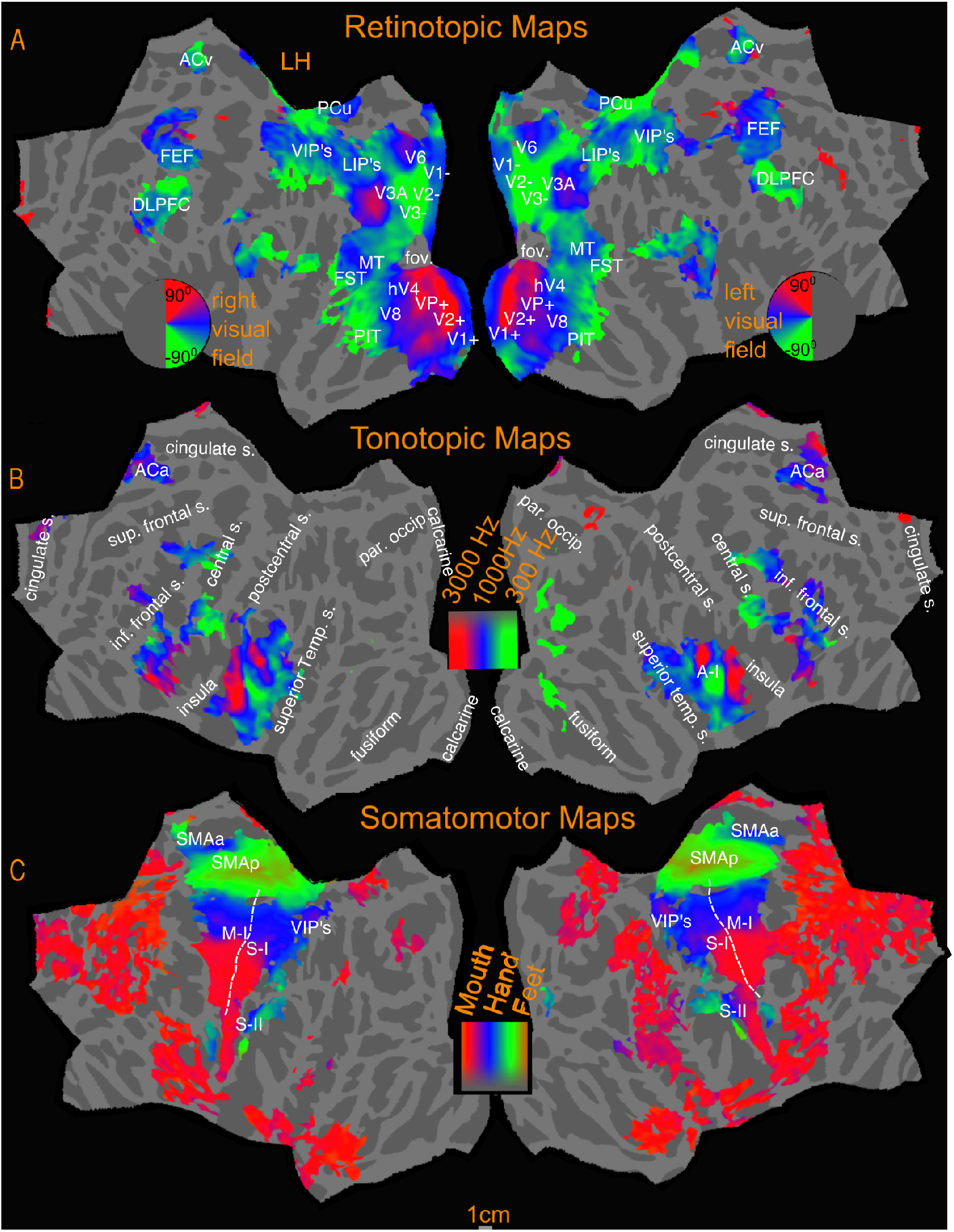
Sensory-motor maps-Polar phase maps. All activations are illustrated at vertex threshold of p<0.05, cluster thresholded to p<0.05

**Figure 7:**
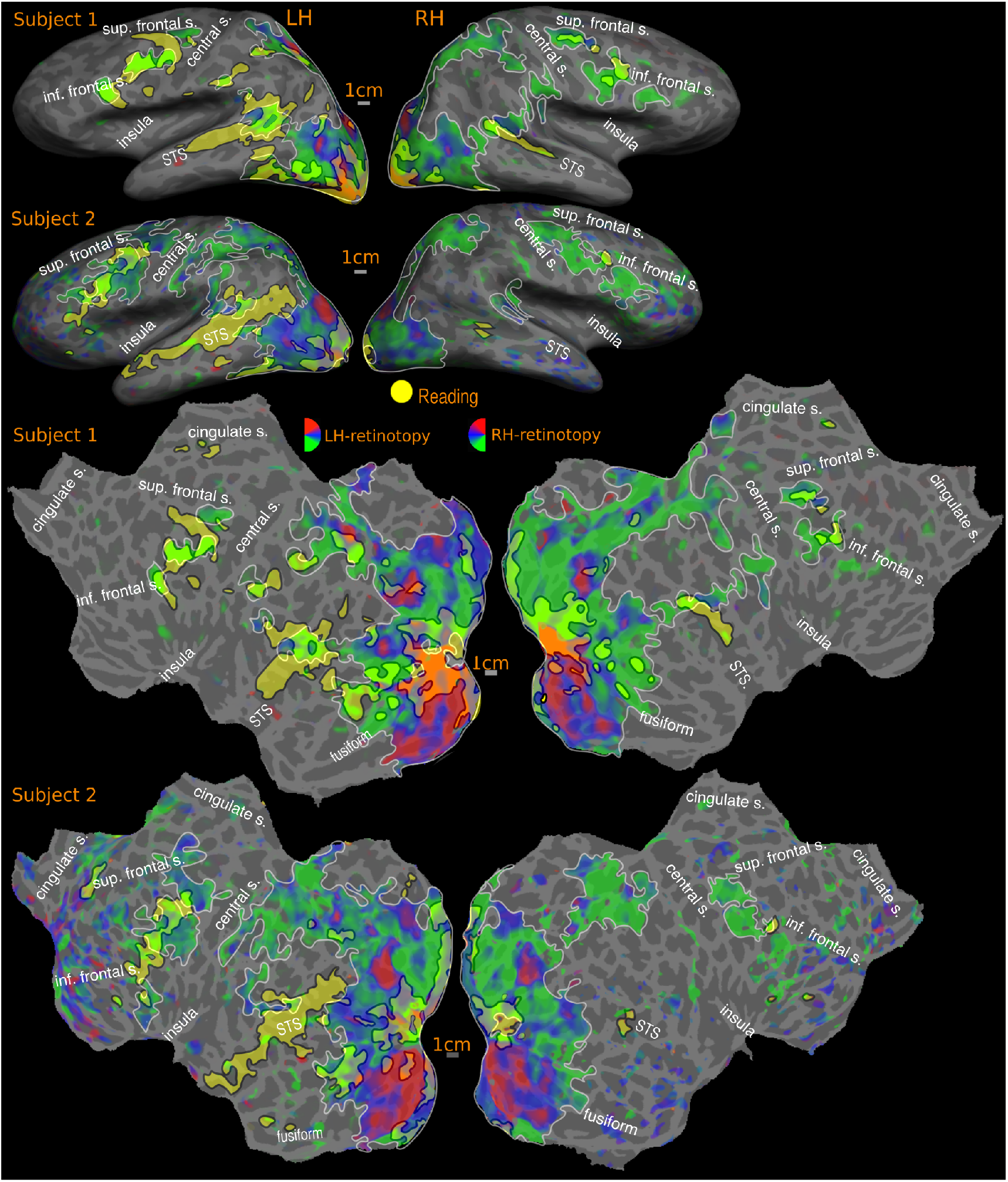
Overlap of READING (English vs. Hindi) with RETINOTOPIC maps-individual SUBJECTS. Activations are illustrated at p<0.001, corrected to p<0.01. Percentage of reading activation overlapping with retinotopy: Subject-1: 58% (LH), 83% (RH); Subject-2: 48% (LH), 81% (RH). Only colors used in the figure are red, blue, green (to represent sensory-motor maps) and yellow (to represent language). Any other colors perceived are the result of overlap of colors (e.g. orange where red and yellow overlaps)

**Figure 8:**
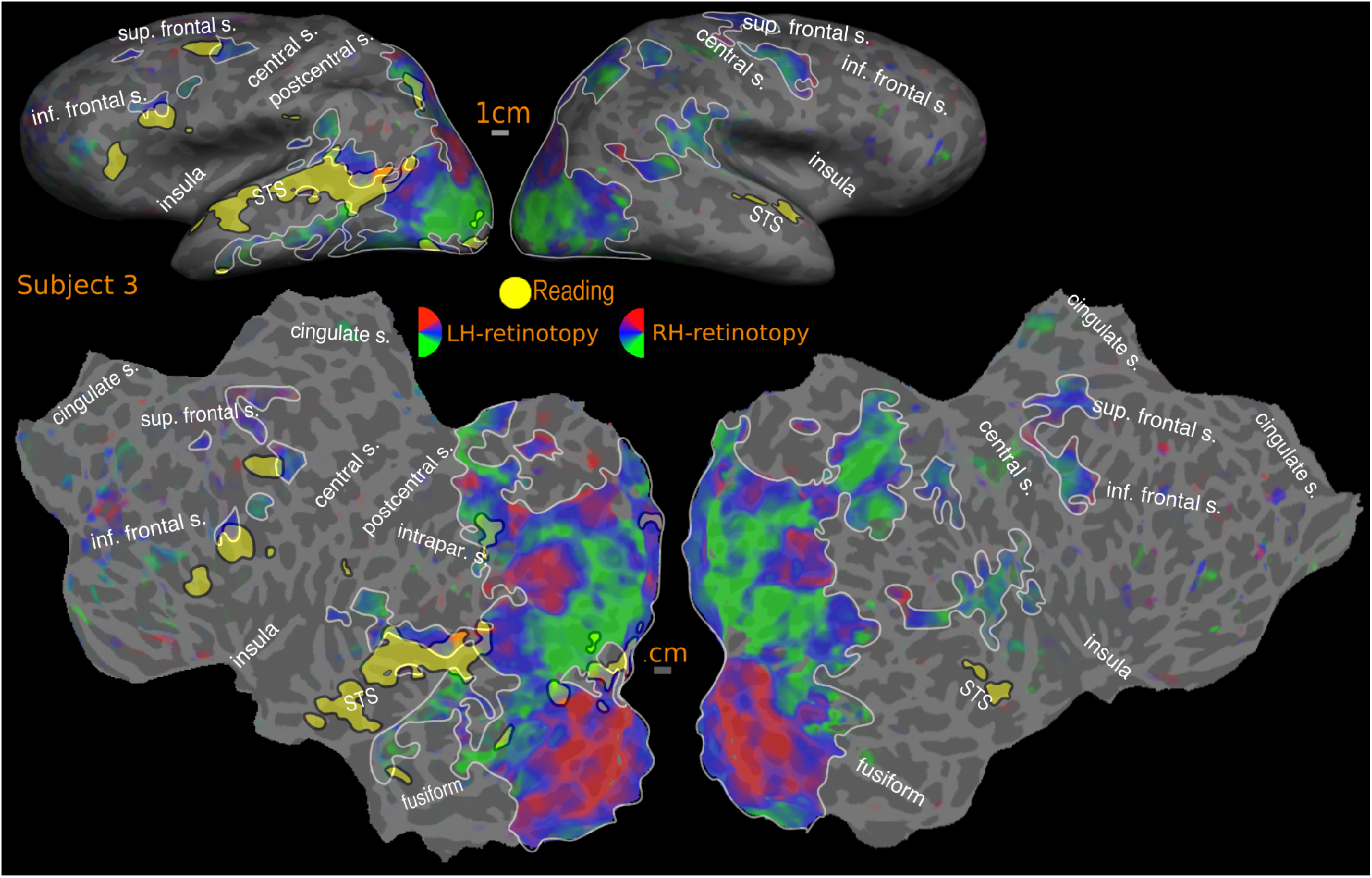
Overlap of READING (English vs. Hindi) with RETINOTOPIC maps-individual SUBJECT. Activations are illustrated at p<0.001, corrected to p<0.01. Percentage of reading activation overlapping with retinotopy: Subject-3: 28% (LH), Nil (RH)

**Figure 9:**
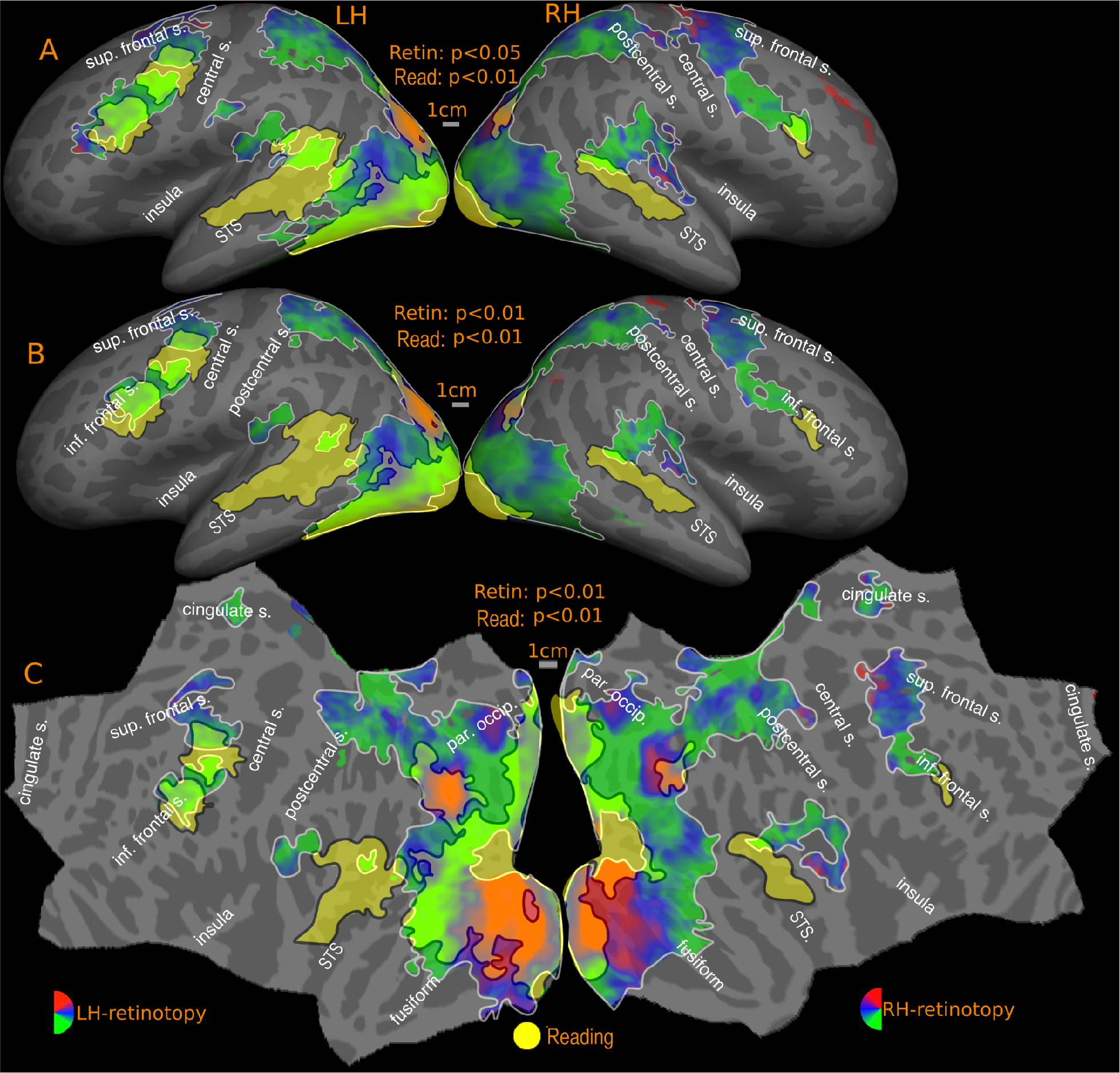
Overlap of READING (English vs. Hindi) with RETINOTOPIC maps-GROUP. Reported p values are vertex thresholds used before cluster thresholding. All activations are cluster thresholded to p<0.05. Reading activation is illustrated at p<0.01 (vertex threshold) in A and B. Retinotopic activation is illustrated at two different vertex thresholds-p<0.05 (A) and p<0.01 (B). C represents the flattened version of B. Percentage of reading activation overlapping with retinotopy-A: 81% (LH), 81% (RH); B: 75% (LH), 73% (RH)

The mapped regions in both lower and higher level visual areas unanimously preferred contralateral visual stimulation. The retinotopic activation pattern in individual subjects (Fig. 7, 8) closely follows the average pattern in Figure 6A. It should be noted that the lack of retinotopy in the result figures around the fovea is an artifact expected with central fixation (no periodic signal change will be present since the subject is fixating at all times); these regions are known to be retinotopic (e.g., Schira et al., 2009) and are considered so for overlap analysis.

### Tonotopic maps

Tonotopy is observed in several regions in the cortex (amplitude in Fig 5C, phase in Fig. 6B; see also phase underlays in Figs. 10, 11). In all phase maps, lower frequencies are green, middle frequencies blue, and high frequencies red. The tonotopic maps in and around primary auditory cortex in the lateral fissure are well known and followed characteristic features previously reported in the literature (e.g., Talavage et al., 2000; 2004; Formisano et al., 2003). In both the group maps and the individual subject maps, there is a high-to-low-to-high frequency progression across Heschl’s gyrus (HG), moving diagonally (medially and anteriorly) across the temporal plane. This region is flanked by higher frequency representation anteromedially towards the planum polare (PP) and posteromedially towards the planum temporale (PT). Posterior to HG on the lateral PT on the superior temporal gyrus, an additional low frequency focus, and two small high frequency foci are also observed. A few subjects also exhibited tonotopic responses extending inferiorly beyond the fundus of the superior temporal gyrus onto the middle temporal gyrus (e.g. subject-3).

Tonotopic maps were also found in several frontal regions (discussed further under New maps). In the cross subject average those regions are most significant in inferior frontal cortex, occupying a region on the precentral gyrus that extends posteriorly into the central sulcus and anteriorly into the precentral sulcus, reaching the inferior/anterior end of the precentral sulcus. While tonotopy in human frontal lobe has received scant attention so far, our results suggest that consistent frontal tonotopy is observed with only a moderate degree of variation across subjects.

Finally, a distinct tonotopic region was found in both hemispheres in the anterior cingulate region (ACa in Fig. 6B, discussed further under New maps). The anterior cingulate tonotopic region is immediately anterior to the anterior cingulate visual maps (ACv) described above, and does not overlap it.

### Somatomotor maps

Figure 12 illustrates the group level somatomotor maps on the inflated surface (also see amplitude, Fig. 5D, and phase maps, Fig. 6C, for a flattened view) overlapped with reading (English vs. Hindi) activation. The somatomotor mapping revealed a highly significant ventral-to-dorsal face-to-leg somatotopic representation in M-I proper and premotor cortex, as well as in a number of post-central areas including primary somatosensory cortex (including areas 3a, 3b, 1, and 2), parts of area 5, two small maps in the upper bank of the lateral sulcus (S-II and related areas), and most of VIP+, where we identified a characteristically medially displaced face representation in the postcentral sulcus (red), as well as a more medial lower-visual-field-overlapping lower body representation areas recently revealed in Huang et al.'s (2012) mapping study that used an air puff body suit. Finally, we activated two additional smaller body representations (leg-to-face and face-to-leg) on the medial surface moving posterior to anterior in supplementary motor area (SMA) at both group & individual level. Though neither of these appears to reach all the way to the face (red) in the average, the reason for this is that inter-subject variation in the exact location of these small maps resulted in flattening the phase range in the average; a full face representation was visible in a number of other individual subjects (data not shown). During mapping of the face, participants softly mouthed syllables, which they were nevertheless able to hear, and which are similar to speech sounds, which we believe explains some face-correlated activation observed in temporal auditory areas. In order to assess the possible auditory contribution, we analyzed the activation due to auditory cue, (by doing a Fourier analysis at the frequency at which auditory cue occurred, 88 cycles/scan and thresholded at p<0.05), and masked out the activated regions. These masked regions mainly involved the auditory cortex and small regions in the insula, frontal cortex and medial anterior cingulate. There is also moderate apparently face-correlated activation in anterior frontal areas and the anterior temporal lobe. This was likely caused by B0 deformations due to changing the distribution of air in the oral cavity during active mouth movements (these brain regions are closest to the oral cavity). There was no map-like structure (phase spread or reversals) in the apparent activity in these regions, nor any overlap with reading activation. By contrast, the region on the anterior bank of the central sulcus near the border between the face and hand representations of M-I – which is substantially further away from the oral cavity – was a poly-sensory zone activated also by passive visual and auditory stimulation, as can be seen in individual maps and group subject maps (Fig. 6).

### Overlap of reading (English vs. Hindi) activation with visual, auditory, and somatomotor maps

Figures 7-13 show the overlap of reading (English vs. Hindi) activation with topological visual, auditory and somatomotor maps. For the cross-subject activations, the visual, auditory, and somatomotor activations were assessed for two different hard vertexwise thresholds before cluster exclusion correction: p<0.01 (higher threshold) and p<0.05 (lower threshold). Reading activation uses a single higher hard vertex threshold of p<0.01. For single subject activations, reading, retinotopic, and tonontopic maps are hard thresholded at p<0.001, corrected to p<0.01. The overlap estimates below, are expressed as the percentage of reading activation intersecting with sensory-motor maps. The quantitative results include an overall estimate, where the percentage of total reading activation overlapping with retinotopic, tonotopic and somatomotor maps is reported for each hemisphere. Additionally, each region (frontal, temporal and occipito-parietal) for reading activation is considered separately, and corresponding overlap is expressed as a percentage of the regional reading activation. The quantitative estimates are summarized in Table 1.

**Figure 10:**
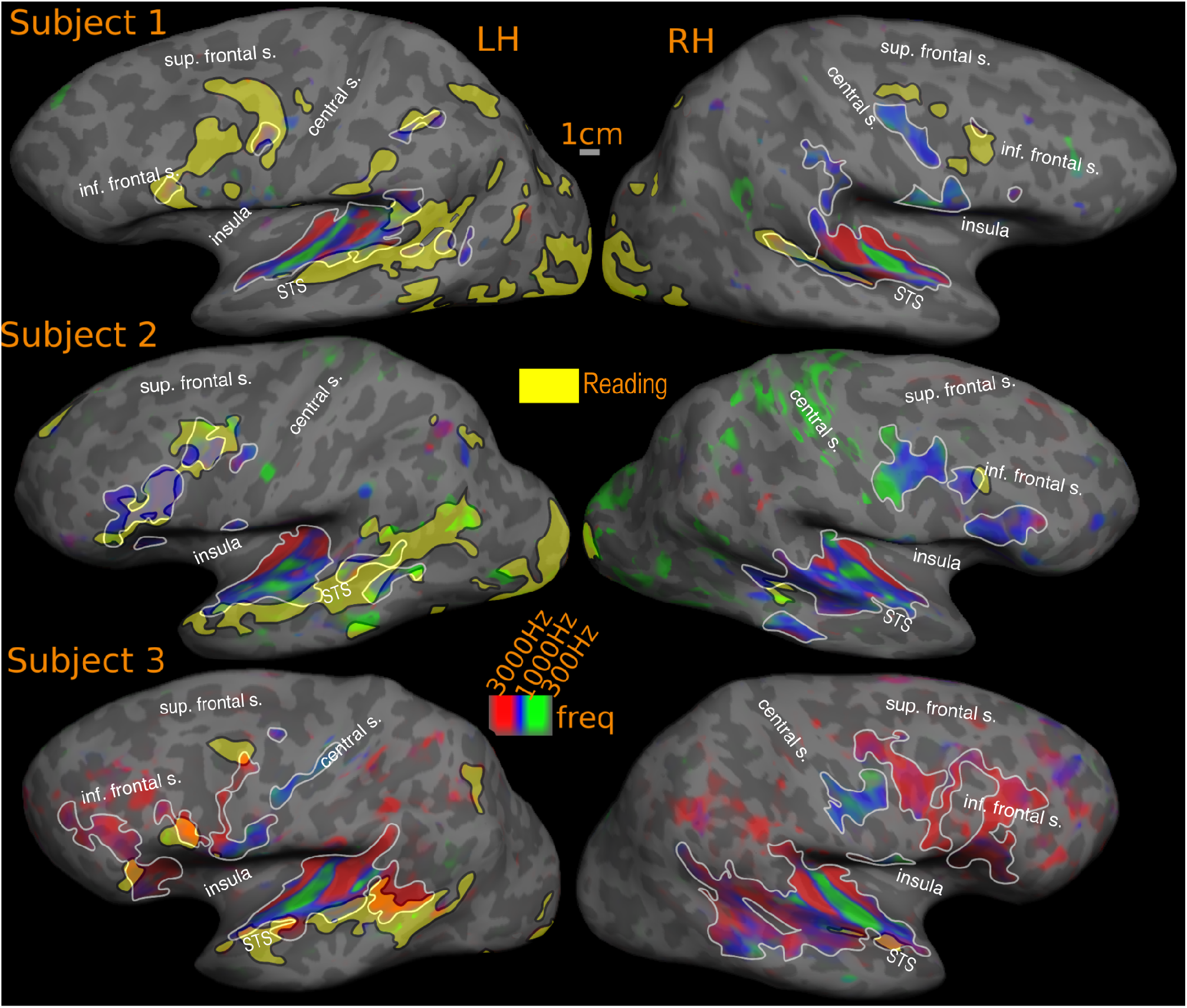
Overlap of READING (English vs. Hindi) with TONOTOPIC maps-individual SUBJECTS. Activations are illustrated at p<0.001, corrected to p<0.01. Percentage of reading activation overlapping with tonotopy: Subject-1: 6% (LH), 10% (RH); Subject-2: 14% (LH), 7% (RH); Subject-3: 32% (LH), 54% (RH)

**Figure 11:**
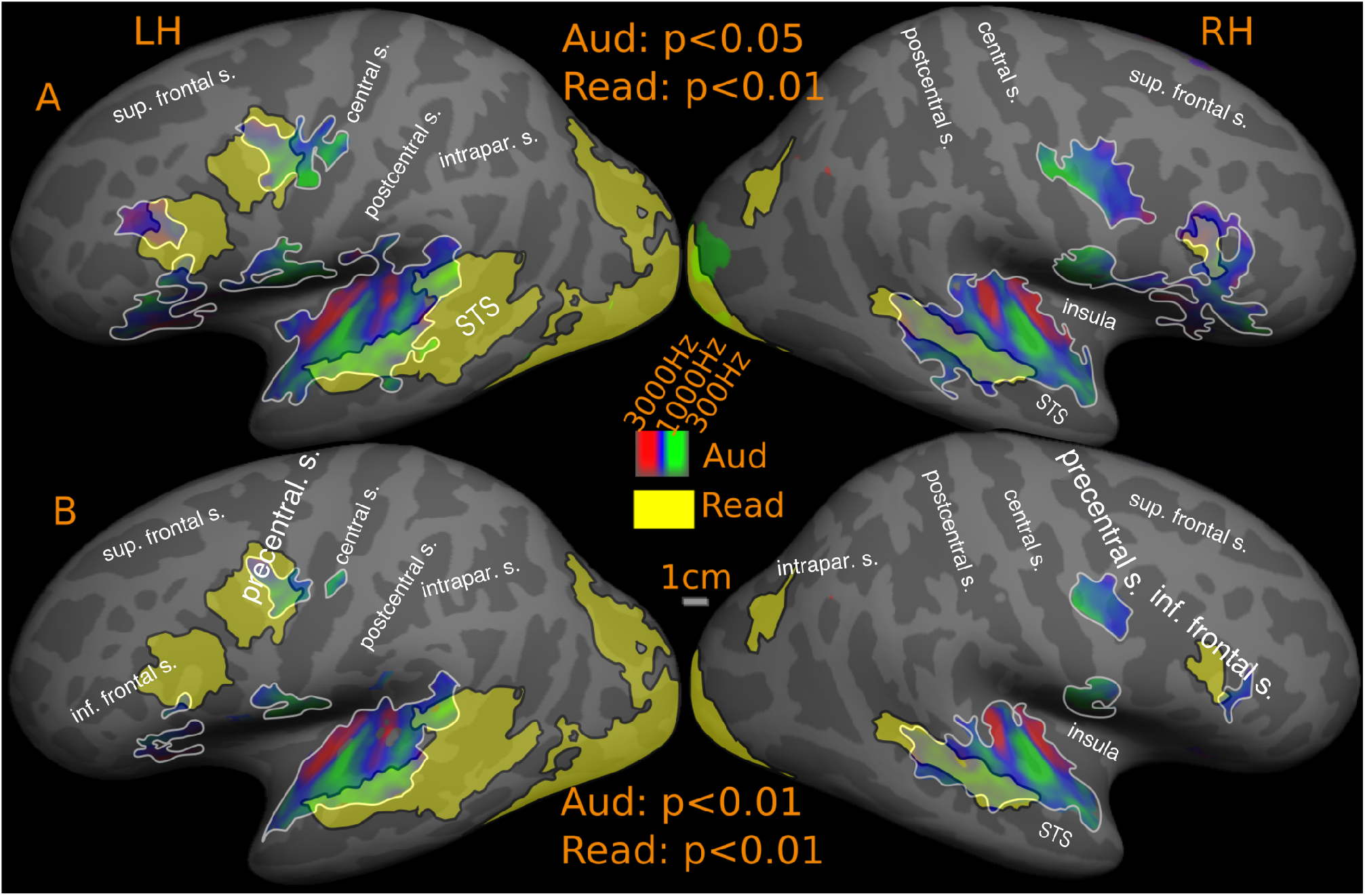
Overlap of READING (English vs. Hindi) with TONOTOPIC maps-GROUP. Reported p values are vertex thresholds used before cluster thresholding. All activations are cluster thresholded to p<0.05. Reading activation is illustrated at p<0.01 (vertex threshold) in A and B. Tonotopic activation is illustrated at two different vertex thresholds-p<0.05 (A) and p<0.01 (B). Percentage of reading activation overlapping with tonotopy-A: 10% (LH), 20% (RH); B: 6% (LH), 13% (RH)

**Figure 12:**
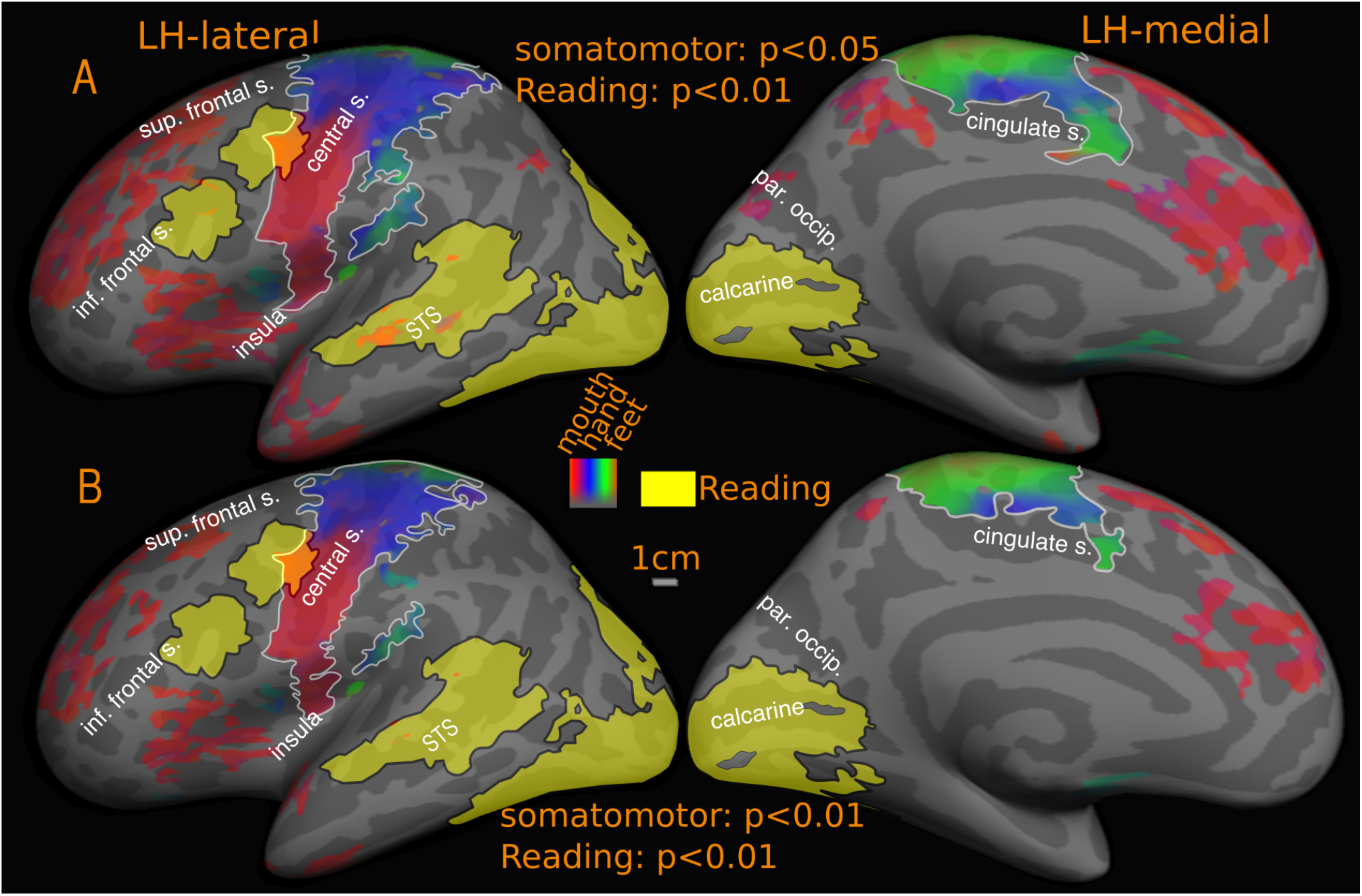
Overlap of READING (English vs. Hindi) with SOMATOMOTOR maps-GROUP. Reported p values are vertex thresholds used before cluster thresholding. All activations are cluster thresholded to p<0.05. Reading activation is illustrated at p<0.01 (vertex threshold) in A and B. Somatomotor activation is illustrated at two different vertex thresholds-p<0.05 (A) and p<0.01 (B). Percentage of reading activation overlapping with somatomotor maps-A: 2% (LH), Nil (RH); B: 1.6% (LH), Nil (RH)

**Figure 13:**
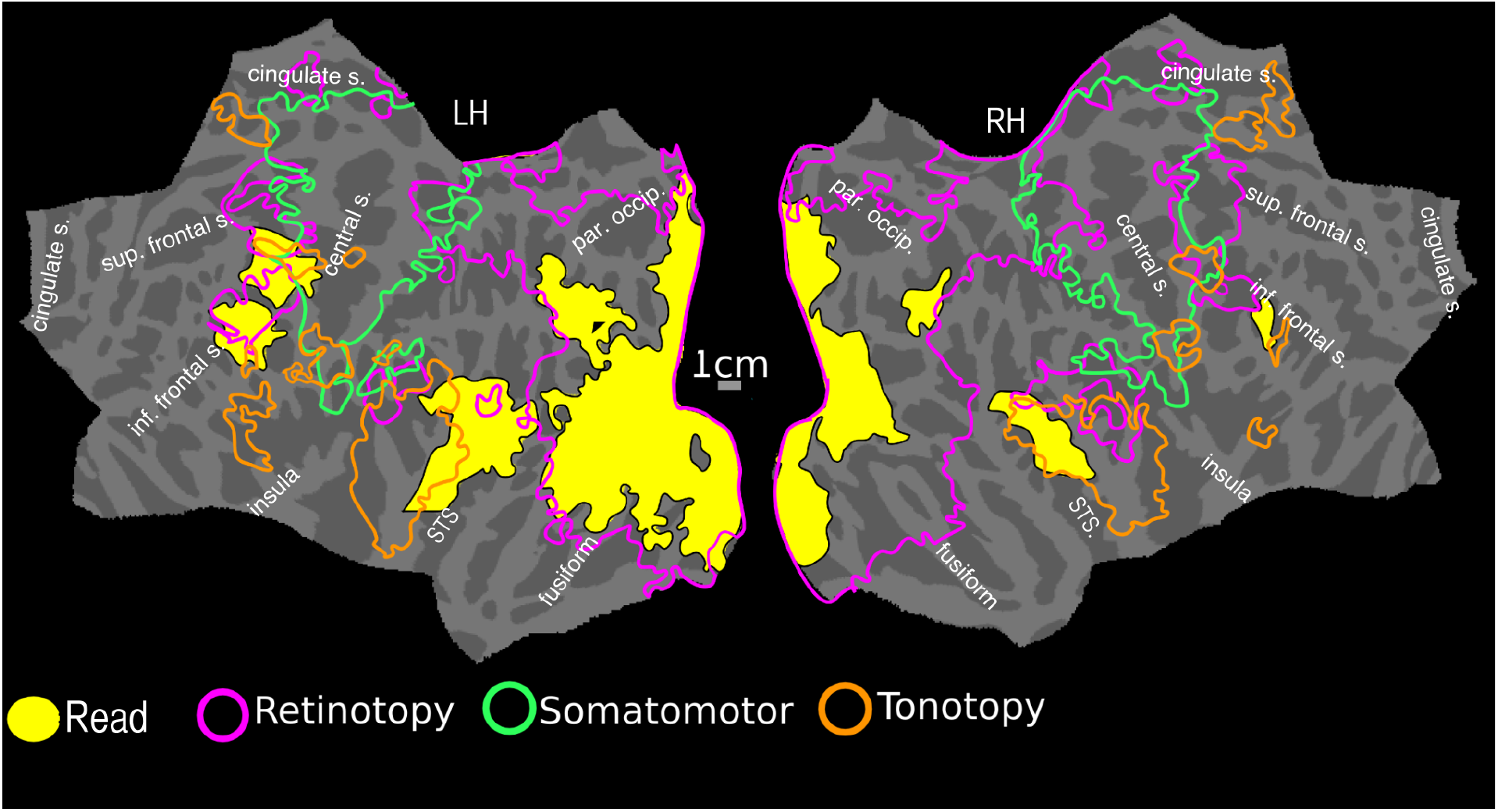
Overlap of READING (English vs. Hindi) with ALL MAPS-GROUP. Reading activation and maps are illustrated at vertex threshold p<0.01, cluster corrected to p<0.05

**Table 1.** Overlap of Reading activation with Retinotopic and Tonotopic maps.

| Retinotopic maps (as a percentage of reading activation) |  |  | Group |  | Subj-1 | Subj-2 | Subj-3 |
| --- | --- | --- | --- | --- | --- | --- | --- |
|  |  |  | Higher threshold | Lower threshold |  |  |  |
|  | Left hemisp here | Overall | 75 | 81 | 58 | 48 | 28 |
|  |  | Frontal | 55 | 70 | 34 | 40 | 11 |
|  |  | Temporal | 7 | 26 | 17 | 5 | 13 |
|  |  | Occipital | 100 | 100 | 100 | 100 | 100 |
|  | Right hemisp here | Overall | 73 | 81 | 83 | 81 | Nil |
|  |  | Frontal | Nil | 62 | 69 | 38 | Nil |
|  |  | Temporal | 8 | 15 | 18 | Nil | Nil |
|  |  | Occipital | 100 | 100 | 100 | 100 | Nil |
| Tonotopic maps (as a percentage of reading activation) | Left hemisp here | Overall | 6 | 10 | 6 | 14 | 32 |
|  |  | Frontal | 10 | 22 | 8 | 51 | 44 |
|  |  | Temporal | 20 | 26 | 17 | 23 | 39 |
|  |  | Occipital | Nil | Nil | Nil | Nil | Nil |
|  | Right hemisp here | Overall | 13 | 20 | 10 | 7 | 54 |
|  |  | Frontal | 6 | 84 | 8 | 33 | Nil |
|  |  | Temporal | 74 | 85 | 71 | 29 | 61 |
|  |  | Occipital | Nil | Nil | Nil | Nil | Nil |

#### Reading overlap with retinotopic maps

The retinotopy/reading overlap for the cross-subject surface average is shown in Figure 9. The reading activation in the whole of occipital and parietal cortex in both hemispheres fall within retinotopically mapped regions. In the left hemisphere, the overlapping regions include early visual areas such as V1, V2, ventral V3, extending to posterior inferior temporal cortex all the way to the lateral surface reaching MT and immediately surrounding areas, after which the lateral activation is attenuated. There is a more discontinuous overlap on the superior bank of the calcarine sulcus, with V1 and V2 activated, but little activation in dorsal V3. There is a strong activation again in the vicinity of V3A and immediately anterior areas. The retinotopic branch extending across MT overlaps with the reading activation in the posterior STS. The rest of the reading activation in the STS is outside the bounds of any retinotopy. In the frontal cortex, there is substantial overlap with both maps in the FEF region as well as more laterally located dorsolateral prefrontal cortex. In the right hemisphere, reading activation in occipital cortex is largely limited to early visual areas V1 and V2. On the lateral occipital side, significant activation is observed immediately anterior to V3A. The STS activation overlaps partially with retinotopic maps in the posterior STS regions, and the right frontal activation overlaps partially with retinotopy in the frontal cortex. The reading activation around the fovea region is considered as overlapping with retinotopy, as the lack of retinotopy in this region is an artifact expected with central fixation (no periodic signal change will be present since the subject is fixating at all times); however, these regions are known to be retinotopic (e.g., Schira et al., 2009) and are considered so for overlap analysis.

With regards to quantitative overlap estimate, over the entire cortex, at the higher threshold, more than 70% of the cross-subject reading activation in both hemispheres falls within retinotopic areas (LH: 75%, RH: 73%); the overlap is more than 80% at the lower threshold (LH: 81%, RH: 81%) (see Fig 9). We previously described three main regions of reading activation – occipito-parietal, superior temporal and frontal. The reading activation in region-1, which includes the occipital lobe, inferior temporal lobe, and superior parietal lobe falls completely within retinotopic regions in both hemispheres (100% overlap). In reading region-2, the superior temporal cortex, in the left hemisphere, the overlaps were 7% (higher threshold) and 26% (lower threshold). In the right hemisphere, the overlaps were 8% (higher threshold) and 15% (lower threshold). For reading region-3, frontal cortex, which includes precentral and inferior frontal cortex, there is a 55% overlap with retinotopy at the higher threshold in the left hemisphere, which rises to 70% at the lower retinotopy threshold. In the right hemisphere, the frontal retinotopic maps lie just outside the borders of right hemisphere reading activation in frontal cortex at the higher threshold; at the lower retinotopy threshold, they extend to cover almost 62% of the right hemisphere frontal reading activation.

For two individual subjects subject-1 and subject-2 (Fig. 7), the overall left hemisphere retinotopy/reading overlap is 53% (subject-1: 58% and subject-2: 48%), while the right hemisphere overlap is above 80% (subject-1: 83% and subject-2: 81%). Zooming in to individual regions, as with the cross-subject average, there is almost 100% overlap between reading and retinotopy in occipito-parietal cortex in both hemispheres. In the left hemisphere frontal and temporal reading areas, overlap with retinotopy is 34% and 17% in subject-1 and 40% and 5% in subject-2. The corresponding overlaps for the right hemisphere of these subjects are 69% (frontal) and 18% (temporal) for subject-1, and 38% (frontal) and 0% (temporal) for subject-2.

In subject-3 (Fig. 8), the frontal and temporal reading profile varies considerably and has a lower overlap (LH: 28%, RH: Nil) with retinotopic maps than it does with tonotopic maps (see below). The individual left hemisphere overlap figures in frontal and temporal cortex are 11% and 13%. There is no overlap in the right hemisphere.

The retinotopic map underlays illustrated in Figures 7-9 use the same color scale as the average (lower contralateral field green, horizontal meridian blue, upper field red). In the group results we found that in both hemispheres, the occipital reading activation overlaps roughly equally with lower, middle, and upper visual fields. In higher-level retinotopic areas, there was a slight predominance of horizontal meridian and lower visual fields. There are two reasons for this. First, because retinotopic maps in higher-level visual areas are smaller, slight inter-subject displacements in the location of these maps has a tendency to reduce the range of polar angles in the average map. This 'regression toward the horizontal meridian' results in an overrepresentation of the horizontal meridian in the average. Another possible source of a horizontal meridian and particularly lower field emphasis is that subjects in our direct-view experiments had to lower their gaze somewhat to fixate the center of the close-up screen. Given that there is evidence for head-centered remapping of receptive fields in some higher visual areas (e.g., see Sereno and Huang, 2006), the lowered gaze may have remapped the entire visual field slightly toward the lower field in these higher level areas, resulting in an overemphasis of the lower field in both individual subjects as well as the average.

#### Reading overlap with tonotopic maps

The most obvious overlap is in temporal cortex, where the reading task consistently activates an anterior-posteriorly elongated region in superior temporal cortex. In the left hemisphere average, the center of the superior temporal reading activation is slightly inferior to the center of the tonotopic activation (Fig. 11, and compare Fig. 5A and 5C) with overlap on the superior temporal gyrus and part of the upper bank of the STS. The inferior edge of the reading activation in turn overlaps the superior edge of middle temporal visual areas. Most subjects (and the average) showed more reading overlap with retinotopic maps than tonotopic maps. However, this was reversed in some subjects (e.g. subject-3). In the right hemisphere average, the smaller superior temporal reading activation almost completely overlaps tonotopy.

The frontal lobe reading activation also overlaps partially with frontal tonotopic maps. These are also potential sites of multi-sensory integration since these region contain both retinotopic and tonotopic maps. These findings are consistent in the cross-subject average as well as in individuals. Though the cluster correction filter removed it as being too small to be valid (and hence it is not formally included in overlap analysis), there is a small but strong reading activation (t>4.7, p<10^-4^) in anterior cingulate, which overlaps almost completely with the tonotopic map in the region. This region is only activated while reading English.

The overall tonotopy/reading overlap (Fig. 11) estimate is 6% (LH) and 13% (RH) for the higher threshold, and 10% (LH) and 20% (RH) for the lower threshold in the cross-subject average. In left temporal cortex, the level of overlap is 20% at the higher threshold and 26% at the lower threshold. In the right hemisphere, more of the temporal reading activation falls within tonotopy (74% at higher threshold, 85% at lower threshold). The tonotopic maps also overlap with frontal reading activation in both hemispheres: 10% (high threshold) and 22% (low threshold) in left hemisphere and 6% (high threshold) and 84% (low threshold) in the right hemisphere.

Our typical individual subjects (Fig. 10) exhibit a similar profile (overall overlap: subject-1: LH: 6%, RH: 10%; subject-2: LH: 14%, RH: 7%). In subject-1, 8% of frontal and 17% of temporal lobe activation in left hemisphere overlaps with tonotopy. The corresponding figures for right hemisphere are 8% frontal and 71% temporal. For subject-2, the left hemisphere overlap is 51% frontal and 23% temporal and for the right hemisphere, 33% frontal and 29% temporal. Subject-3 has especially extensive tonotopic maps and shows a higher degree of overall overlap with tonotopy than with retinotopic areas (overall: 32% in left hemisphere and 54% in right hemisphere). Around 44% of frontal and 39% of temporal lobe left hemisphere activation falls within tonotopically defined regions. In the right hemisphere, nearly 61% of temporal reading activation overlaps with tonotopy. No significant frontal reading activation is observed for subject-3 in the right hemisphere; and there are tonotopic maps in this region.

In the group results, the majority of the overlap between reading and tonotopy in the temporal lobe falls within the low to middle frequency range. For example, in the left STG, only 0.7% of temporal reading activation overlaps high frequencies. In the right temporal cortex, reading/tonotopy overlaps occur within the mid-low frequency range avoiding high frequency regions in the lateral sulcus. The situation in frontal cortex is more mixed. In the left frontal cortex, the overlapping region lying between precentral and central sulcus is dominated by mid-low frequencies closer to central sulcus and by high-mid frequencies closer to the precentral sulcus. In the pars opercularis region, there are two separate overlaps, the more dorsal overlap region is predominantly high-mid range frequencies while the more ventral region is predominantly mid range frequencies. In the right hemisphere, the overlap is only with the pars opercularis activation, where the tonotopic maps overlapping with reading has a high-mid frequency distribution.

#### Reading overlap with somatomotor maps

In the cross-subject average, there is only one overlap zone, near the face/hand boundary bordering the central sulcus in M-I (Fig. 12). The overlapping region corresponds to 1.6% of total reading activation at higher threshold. This constitutes 12% (high threshold) of the left frontal reading activation, rising to 14% at low threshold. There was no overlap in right hemisphere or SMA.

### New retinotopic and tonotopic maps

The topological mapping experiments uncovered several sensory maps previously unreported in the literature. In nonhuman primates, it has long been known that the anterior cingulate supplementary motor areas are directly adjoined by the dorsomedial frontal eye fields in primates (Schiller and Chou, 1998; Purcell et al., 2012). So far, there have not been any reports on topological maps in a similar region in humans. Our data may provide the first observation of the topological organization of the human equivalent of the dorsomedial frontal eye fields (ACv in Figure 6A), which posses a retinotopic map of the contralateral visual field (their location is more inferior than in non-human primates, in line with similar superior-toward-medial-wall movements of parietal retinotopic areas in humans comparable to LIP). While retinotopy in this region is reported here for the first time, there have been a number of reports of activation in this region associated with visuo-spatial attention and eye movements (Mesulam et al., 2001; Pierrot-Deseilligny et al., 2004; McDowell et al., 2008; Jamadar et al., 2013; O’Reilly et al., 2013). In the reading experiment, all three conditions that involved eye movements-English, Hindi and Dot have significant activation overlapping the anterior cingulate retinotopic region. This activation disappeared in the English-Hindi and Hind-Dot contrasts suggesting the activation is more relevant for eye movements or/and spatial attention than for comprehension per se. Adjacent to the dorsomedial retinotopic map, we also found a previously unreported tonotopic map, here after referred to as the dorsomedial frontal ‘ear’ fields (ACa in Figure 6B). The anterior cingulate retinotopic map and the tonotopic map are separated by SMA map and do not overlap each other or SMA. There is strong reading activation aligned with the tonotopic map in this region and unlike the reading activation in the anterior cingulate retinotopic region, this activation is strongest while reading English, compared to both the other conditions. The relatively smaller activation observed in Hindi and Dot disappears in the Hindi-Dot contrast, while English-Hindi has a significant activation, in alignment with dorsomedial frontal ‘ear’ fields (Figure 4 and Figure 5A. Not shown in overlap figures, as cluster correction algorithm excluded this small region). Finally, we found several previously unreported tonotopic maps in frontal cortex. The frontal tonotopy partially overlaps with retinotopic maps in the region as well as the reading activation found in the frontal cortex.

## Discussion

The results presented here provide a comprehensive qualitative and quantitative assessment of where, and by how much, processes relevant to naturalistic reading overlap with topological (neighbor-preserving) visual, auditory, and somatomotor maps using high-resolution surface-based fMRI across the entire cortex. Nearly 80 fMRI sessions were analyzed with a start-to-finish cortical-surface-based processing pipeline. This study combines recent advances in lower-level sensory-motor mapping with a higher-level cognitive task (naturalistic reading with matched eye movements across conditions). There were two related objectives.

The first was to accurately localize regions of interest in naturalistic reading comprehension by determining their exact relation to low and high-level topological visual, auditory, somatosensory and motor maps. Topological mapping is a time-tested method for accurately defining the boundaries of cortical areas. The figures presented here are the first illustrations of the relative location of naturalistic reading comprehension activation and topological visual, auditory and somatomotor maps across the entire cortex in the same group of subjects. In the process, we discovered several previously unreported tonotopic maps in frontal cortex that partially overlapped frontal retinotopic maps. In anterior cingulate cortex, we found two additional new maps – a visual map and a tonotopic map, both near the SMA but not overlapping each other, or the SMA. A strong reading (English vs. Hindi) activation overlapped the left anterior cingulate tonotopic map. Our data sets benefitted from the latest multi-band pulse sequences, allowing us to map the whole brain without losing temporal SNR, as well as from optimized sensory mapping stimuli, which together may account for why we were able to visualize those new maps.

Second, we wanted to provide a quantitative estimate for the level of overlap between activations observed during naturalistic reading comprehension and topological sensory-motor maps, which are driven by relatively low-level, sensory-motor stimuli. Our finding is that 80% of reading activations (English vs. Hindi) in the left hemisphere and 86% in the right hemisphere fall within regions containing topological visual, auditory, somatosensory, or motor maps in the group average. At least 65% of the reading activation in frontal cortex, 26% in temporal cortex, and 100% in parietal and occipital cortex falls within visual, auditory or somatomotor maps. The overlap figures rise to nearly 90% in frontal cortex and 50% in temporal cortex if the threshold for detecting topological mapping is lowered to p<0.05 (corrected). Given that smaller, higher-level maps are more difficult to visualize than large, metabolically active, early visual areas, these figures serve as a conservative rather than an overly optimistic estimate (see Gonzales-Castillo et al., 2012). We should also point out that the 'overlap' defined here could only be measured at the resolution of our fMRI voxels. An overlap voxel might be identified if the two kinds of signals were within the same neuron, within distinct but adjacent neurons, within different 50 micron wide minicolumns, or within different ∽1mm wide columns (see e.g., Lund et al., 1993), all of which could be within the same voxel.

While occipital cortex and immediately surrounding regions have long been considered to be quintessentially visual, language research has often regarded activation elsewhere as falling outside the bounds of vision. For example, the classical language areas in superior temporal cortex and frontal cortex are often considered to be non-visual. However, in the last decade or so, several additional retinotopic maps were discovered in frontal cortex (Hagler and Sereno, 2006; Hagler et al., 2007; Kastner et al., 2007), parietal cortex (Sereno et al., 2001, 2003; Huk et al., 2002; Schluppeck et al., 2005; Silver et al., 2005; Sereno and Huang, 2006; Swisher et al., 2007; Silver and Kastner, 2009) and temporal cortex (Sereno et al., 2003; Huang and Sereno, 2013). The higher-level retinotopic maps in frontal cortex were originally discovered using working memory and spatial attention paradigms (Hagler and Sereno, 2006; Kastner et al., 2007). Those studies suggested that topological maps may serve as a convenient method of allocating working memory, or maintaining pointers to specific content, even (or more so!) for tasks that don't explicitly require attention to spatial location, but that require attention to content. The present study shows that reading activation in frontal cortex falls largely within these retinotopic maps.

In addition to confirming previous reports of retinotopic maps in frontal cortex, we also found new evidence for consistent tonotopy both in dorsolateral prefrontal cortex and ventral prefrontal cortex. While tonotopy in superior temporal cortex has been the subject of several studies (Talavage et al., 2000, 2004; Formisano et al., 2003; Humphries et al., 2010; Da Costa et al., 2011; Dick et al., 2012; Langers and Van Dijk, 2012) there is virtually no literature on tonotopy in human frontal cortex. There is, by contrast, a wealth of neuroimaging studies providing evidence that the human frontal lobe is active during general auditory tasks (Platel et al., 1997; Alain et al., 2001; Kiehl et al., 2001; Muller et al., 2001; Gaab et al., 2003; Arnott et al., 2004; Rämä et al., 2004; Koelsch et al., 2009). Neurophysiological and neuroanatomical findings in non-human primates show that both dorsolateral and ventral prefrontal cortex is reciprocally interconnected with auditory regions in the temporal cortex (Hackett et al., 1999; Romanski et al., 1999; Plakke and Romanski, 2014). The frontal reading activation overlaps the tonotopic maps in the region, perhaps for a similar reason as for the retinotopic map overlap (e.g., memory allocation using topological maps as pointer buffers).

Any activation of higher-level visual and auditory areas during reading comprehension has often been dismissed with the claim that readable, understandable text (here English) attracts more attention than unreadable, incomprehensible text (here Hindi characters of the same word length, fixated with the same controlled pattern of saccades). An assumption prevalent in the literature is that comprehension and reading processes are dissociable from sustained attention, and that they can be dissociated by using a control task matched in attentional complexity. Recent research however suggests that attention is a cortex-wide dynamic process whereby resources are flexibly allocated to the attended function as opposed to a simple mechanism that merely modulates baseline response level (Çukur et al., 2013, Peelen & Kastner, 2014). But this also implies that subtractive analyses attempting to control for attention factors assuming a static modular nature of attention may be overly conservative. In the present study, any task-related attention and working memory processes are considered relevant and necessary for naturalistic reading comprehension and have not been subtracted out. Low level visual processing including saccade preparation and execution (eye movements), and task-irrelevant attention are controlled by subtracting out activation due to the combination of ‘Hindi’ font, directed naturalistic saccades, and the target detection task. In addition to facilitating a naturalistic reading experience, the presentation of text by highlighting the next word drives the exogenous attentional network (Posner, 1980) and triggers a saccade towards the highlighted spatial location. While spatial attention can be modulated without saccades, saccades rarely occur without being directed by attention. Substantial evidence exists suggesting that an eventual saccade target peripheral to fixation attracts attention during saccadic planning (Melcher & Colby, 2008; Zhao et al., 2012), and that attention then quickly returns to the centre of gaze after the saccade to the target has been made. Finally, the random colour change detection task performed by the subjects during the experiment across all conditions served as a control for endogenous attention. In all three conditions-English, Hindi and Dot, significant activation was observed bilaterally near intra parietal/postcentral sulcus, FEF, and dorsomedial frontal ‘eye’ fields, regions known to be activated by visuo-spatial attention and eye saccades (Pierrot-Deseilligny et al., 2004; McDowell et al., 2008; Müri, and Nyffeler, 2008; Jamadar et al., 2013; O’Reilly et al., 2013). The activations in dorsomedial frontal ‘eye’ fields and parts of parietal cortex are fully attenuated in English vs. Hindi contrast (as well as in Hindi vs. Dot contrast, in fact Hindi and Dot has slightly better activations in these areas). In FEF, while the activation disappears in Hindi vs. Dot contrast, a good part of it is still highly significant in English vs. Hindi contrast, suggesting that the FEF may have a larger role in comprehension, an observation supported by other studies (Hasson et al., 2008; Choi et al., 2014).

However, not all frontal regions were activated for all three conditions. The more anterior activation near left inferior frontal sulcus/gyrus (a classical language region) overlapping with the DLPFC visual maps and tonotopic maps were unique to reading and comprehending English passages. Similarly the activation observed in left cingulate dorsomedial tonotopic maps in those regions.

As mentioned above, part of the activation observed could be due to sustained attention and working memory processes engaged in reading meaningful passages; but the functional dissociation of reading processes was very specifically not the goal of the current study. Reading in real situations is a computationally intense, attention- and working memory-demanding contextual process that involves tying together information gleaned from linear symbol streams across seconds and minutes. The idea that these processes are recruited in language processing is not a new one (see Duncan et al., 2000). What is not clear though from the present literature is where the boundaries of these putative multi-functional regions or classical frontal language regions lie. Although we cannot distinguish between domain general processes and purely linguistic processing from the contrast used here, the present study does define precisely where reading activation zones are relative to topological cortical maps in the frontal cortex. Dissociating linguistic and non-linguistic elements in the frontal activation observed in the English vs. Hindi contrast is an important question for future research and we believe similar contrasts using non-linguistic stimuli will help delineate functional specifications in this region, without downplaying comprehension specific processing taking place in these regions.

Despite being a silent reading task, there is also significant overlap with low-level tonotopic maps in the superior temporal gyrus. It has been reported previously that tonotopic maps in auditory cortex overlap with temporal voice areas (Belin, 2000) in STG (Moerel et al., 2012). Additionally, silent active reading has been shown to activate temporal voice areas (Perrone-Bertolloti et al., 2012). Our results are consistent with these findings. Our results also confirm a previous finding that the tonotopic maps overlapping reading activation in the STG have a bias towards the lower frequencies (Moerel et al., 2012), a range occupied by human voices.

There are regions of reading activation, notably in superior temporal cortex, that do not overlap with topological maps; and there are similar, though smaller, non-overlapping regions in frontal cortex near the pars opercularis region. Whether or not these regions are specialized for purely linguistic processing is an important question for future research. Some recent evidence suggests that these regions may have specific functions beyond language processing (see e.g., Dick et al., 2011).

We found increased activation specific to reading meaningful paragraphs in occipital cortex and adjacent regions in inferior/middle temporal lobes and in the fusiform gyrus. Although these regions are not often considered as significant for high level language processing, empirical evidence from lesion studies (Rubens and Kertesz, 1983) and invasive electrophysiology (Burnstine et al., 1990; Luders et al., 1991; Krauss et al., 1996; Mani et al., 2008) suggests that the 'basal temporal language area' in the inferior temporal lobe and fusiform gyrus is as strongly linked with language functions as classical language areas. Electrical stimulation of these regions not only interferes with reading and language understanding but also often leads to speech arrest. Judging from published reports, this area may overlap some of our inferotemporal retinotopy. In addition, in the aphasiology literature, a prominent aphasia called the transcortical sensory aphasia, where the patient exhibits poor ‘Wernicke’-like comprehension, is associated with brain lesions most often in inferior temporal cortex, sometimes including the basal temporal language area, but commonly also involving visual areas in the lateral aspect of the adjoining occipital lobe (Kertesz et al., 1982; Sharp et al., 2004). It is interesting in this context that picture ‘lexical items’ inserted into text sentences can be effortlessly integrated into ongoing linguistic discourse comprehension, even at faster-than-normal reading speeds (e.g., Potter et al., 1986). Our results show that the regions activated during text comprehension in occipital cortex encompass a number of distinct visual areas, both low level (such as V1 and V2) as well as higher level (such as ventral V3, MT, V8, posterior inferotemporal cortex, and LIP+). Taken together, this psychological, brain lesion, and neuroimaging data suggests that some aspects of linguistic meaning assembly may be taking place within intermediate level visual areas. It should be noted that not all activation observed in the occipital cortex for English vs. Fixation was significant for English vs. Hindi contrast. It was instructive to find out where these activations (for English vs. Hindi) lie relative to well-defined distinct retiniotopic maps in posterior occipital cortex.

In contrast to the substantial overlap with visual and auditory maps, somewhat surprisingly, we found rather less overlap of reading activations with somatomotor maps. We did not attempt to select semantic content in our reading task specific for any specific sensory modality and used simple informative passages dealing with natural phenomena, science, information about famous people and paintings, and so on; this may have resulted in less somatosensory content than visual and auditory content. A future direction would be to see if modality specific content modulated the amount of measured overlap with particular modality-specific topological maps. For somatomotor maps, the primary overlap zone was a frontal multimodal overlap zone adjoining primary motor cortex (see next).

There were multisensory maps in several locations including lateral prefrontal and in the extreme superior and posterior part of the lateral sulcus near the supramarginal gyrus. The frontal multisensory overlap zone also overlapped reading activation, but the posterior lateral sulcus zone did not. Note that we have defined 'multisensory' in the restricted sense of voxels that contain topological maps in more than one modality (not voxels that merely respond to more than one modality).

The pattern of activations for the reading experiment across the whole cortex is similar to what was found in a recent natural reading experiment reported by Choi et al. (2014). Compared to Choi et al., our study utilized controlled/ matched saccades as opposed to free eye movements, and a higher resolution, fully surface-based analysis stream as opposed to a volume-based cross-subject analysis only mapped to an average surface at the end. As with our study, the activation patterns in Choi et al. exhibited eye movement related activation patterns in IPS, the FEF, and the dorsomedial frontal eye fields, as well as unique activation for text in more anterior IFG. As observed in our study, Choi et al. also report no activations during natural reading in somatosensory/motor cortex. Visual activation – likely within retinotopic areas judging from our data – was observed even when comparing text with pseudo-words – a closer comparison than ones we used.

There is often an implicit assumption that whatever is going on in higher level visual areas can only count as being involved in language comprehension if it were equally activated by an auditory language understanding task – that is, by this definition, there can be no modality-specific involvement in language comprehension. Given how poorly understood the cortical neural activity patterns underlying language comprehension currently are, we think it is important to keep an open mind about modality specificity or functional duplication. It seems quite possible that, for example, anaphor processing may employ modality-specific representations; the cortical computations and cortical areas involved in dereferencing an auditory “that” could well be different than for resolving a printed “that” that appeared in a particular location on the page. It is difficult to directly experimentally address the question of modality-specific language activations. For example, simultaneous, retinotopically overlapping visual tasks, widespread functional disruption of higher visual areas (e.g., by transcranial magnetic stimulation), or widespread lesions to higher visual areas would disrupt naturalistic reading comprehension itself – the object of study.

Overall, our results show that topological sensory-motor maps, which play crucial functional roles in primary sensory and motor areas, are also found in some of the brain regions involved in a higher-level cognitive function such as reading comprehension. This study provides the first comprehensive overview on the interface between a high-level complex cognitive process reading comprehension, and topologically organized visual, auditory, and somatomotor representations. Determining the specific roles played by the regions identified here remains an important question for future research in the field of topological sensory-motor maps as well as language.

## Acknowledgments

We thank Dr. Frederic Dick for the auditory mapping stimulus.

## References

Alain C, Arnott SR, Hevenor S, Graham S, Grady CL (2001): “What” and “where” in the human auditory system. Proc. Natl. Acad. Sci. U.S.A. 98:12301–12306.

Arnott SR, Binns MA, Grady CL, Alain C (2004): Assessing the auditory dual-pathway model in humans. Neuroimage 22:401–408.

Bandettini PA, Jesmanowicz A, Wong EC, Hyde JS (1992): Processing strategies for time course data sets in functional MRI of the human brain. Magnetic Resonance Medicine 30:161–173.

Burnstine TH, Lesser RP, Hart J, Uematsu S, Zinreich SJ, Krauss GL, Fisher RS, Vining EPG, Gordon B (1990): Characterization of the basal temporal language area in patients with left temporal lobe epilepsy. Neurology 40:966–970.

Belin P, Fillion-Bilodeau S, Gosselin F (2008): The Montreal Affective Voices: a validated set of nonverbal affect bursts for research on auditory affective processing. Behavior research methods 40:531–539.

Belin P, Zatorre RJ, Lafaille P, Ahad P, Pike B (2000): Voice-selective areas in human auditory cortex. Nature 403:309–312.

Choi W, Desai RH, Henderson JM (2014): The neural substrates of natural reading: a comparison of normal and nonword text using eyetracking and fMRI. Frontiers in human neuroscience, 8:1–11. Article 1024.

Cox RW (2012): AFNI: what a long strange trip it’s been. Neuroimage 62:743–747.

Çukur T, Nishimoto S, Huth AG, Gallant JL (2013): Attention during natural vision warps semantic representation across the human brain. Nature neuroscience 16:763–770.

Da Costa S, van der Zwaag W, Marques JP, Frackowiak RSJ, Clarke S, Saenz M (2011): Human primary auditory cortex follows the shape of Heschl’s gyrus. Journal of Neuroscience 31:14067–14075.

Dick F, Lee HL, Nusbaum H, Price CJ (2011): Auditory-motor expertise alters "speech selectivity" in professional musicians and actors. Cerebral Cortex 21:938–948.

Dick F, Tierney AT, Lutti A, Josephs O, Sereno MI, Weiskopf N (2012): In vivo functional and myeloarchitectonic mapping of human primary auditory areas. Journal of Neuroscience 32:16095–16105.

Duncan J, Seitz RJ, Kolodny J, Bor D, Herzog DH, Ahmed A, Newell FN, Emslie H (2000): A Neural Basis for General Intelligence. Science 289:457–460.

Engel SA, Glover GH, Wandell BA (1997): Retinotopic organization in human visual cortex and the spatial precision of functional MRI. Cerebral Cortex 7:181–192

Engel SA, Rumelhart DE, Wandell BA, Lee AT, Glover GH, Chichilnisky EJ, Shadlen MN (1994): fMRI of human visual cortex. Nature 369:525–525.

Ercsey-Ravasz M, Markov NT, Lamy C, Van Essen DC, Knoblauch K, Toroczkai Z, Kennedy H (2013): A predictive network model of cerebral cortical connectivity based on a distance rule. Neuron 80:184–197.

Felleman DJ, Van Essen DC (1991): Distributed hierarchical processing in the primate cerebral cortex. Cerebral Cortex 1:1–47.

Fischl B, Sereno MI, Dale AM (1999a): Cortical Surface-Based Analysis. II. Inflation, flattening, and a surface-based coordinate system. Neuroimage 9:195–207.

Fischl B, Sereno MI, Tootell RB, Dale AM (1999b): High-resolution intersubject averaging and a coordinate system for the cortical surface. Human brain mapping 8:272–284.

Formisano E, Kim DS, Di Salle F, Van de Moortele PF, Ugurbil K, Goebel R (2003): Mirror-symmetric tonotopic maps in human primary auditory cortex. Neuron 40:859–869.

Franceschini S, Gori S, Ruffino M, Pedrolli K, Facoetti A (2012): A causal link between visual spatial attention and reading acquisition. Current biology 22(9):814–819.

Gaab N, Gaser C, Zaehle T, Jancke L, Schlaug G (2003): Functional anatomy of pitch memory—an fMRI study with sparse temporal sampling. Neuroimage 19:1417–1426.

Gabrieli JDE, Norton ES (2012): Reading abilities: importance of visual-spatial attention. Current biology 22(9):298–299.

Gonzalez-Castillo J, Saad ZS, Handwerker DA, Inati SJ, Brenowitz N, Bandettini PA (2012): Whole-brain, time-locked activation with simple tasks revealed using massive averaging and model-free analysis. Proc. Natl. Acad. Sci 109:5487–5492.

Greve DN, Fischl B (2010): Accurate and robust brain image alignment using boundary based Registration. Neuroimage 48:63–72.

Hackett TA, Stepniewska I, Kaas JH (1999): Prefrontal connections of the parabelt auditory cortex in macaque monkeys. Brain Research 817:45–58.

Hagler DJ, Riecke L, Sereno MI (2007): Parietal and superior frontal visuospatial maps activated by pointing and saccades. NeuroImage 35:1562–77.

Hagler DJ Jr, Sereno MI (2006): Spatial maps in frontal and prefrontal cortex. Neuroimage 29:567–577.

Hasson U, Yang E, Vallines I, Heeger DJ, Rubin N (2008): A hierarchy of temporal receptive windows in human cortex. The Journal of neuroscience 28(10):2539–50.

Hebb D (1949): The Organization of Behavior: A Neuropsychological Theory. New York (NY). John Wiley.

Huang R-S, Chen C, Tran AT, Holstein KL, Sereno MI (2012): Mapping multisensory parietal face and body areas in humans. Proc. Natl. Acad. Sci. U.S.A. 109:18114–198119.

Huang R-S, Sereno MI (2013): Bottom-up Retinotopic Organization Supports Top-down Mental Imagery. The open neuroimaging journal 7:58–67.

Huk AC, Dougherty RF, Heeger DJ (2002): Retinotopy and functional subdivision of human areas MT and MST. Journal of Neuroscience 22:7195–7205.

Humphries C, Liebenthal E, Binder JR (2010): Tonotopic organization of human auditory cortex. Neuroimage 50:1202–1211.

Jamadar SD, Fielding J, Egan GF (2013): Quantitative meta-analysis of fMRI and PET studies reveals consistent activation in fronto-striatal-parietal regions and cerebellum during antisaccades and prosaccades. Front. Psychol 4:749.

Jenkinson M, Bannister PR, Brady JM, Smith SM (2002): Improved optimisation for the robust and accurate linear registration and motion correction of brain images. NeuroImage 17:825–841.

Jenkinson M, Smith SM (2001): A global optimisation method for robust affine registration of brain images. Medical Image Analysis 5:143–156.

Kaas JH (1997): Topographic maps are fundamental to sensory processing. Brain Res. Bull 44:107–112.

Kastner S, DeSimone K, Konen CS, Szczepanski SM, Weiner KS, Schneider KA (2007): Topographic maps in human frontal cortex revealed in memory-guided saccade and spatial working-memory tasks. Journal of Neurophysiology 97:3494–3507.

Kertesz A, Sheppard A, MacKenzie R (1982): Localization in transcortical sensory aphasia. Arch Neurol 39:475–478.

Kevan A, Pammer K (2008): Making the link between dorsal stream sensitivity and reading 19(4):67–70.

Kiehl KA, Laurens KR, Duty TL, Forster BB, Liddle PF (2001): Neural sources involved in auditory target detection and novelty processing: an event-related fMRI study. Psychophysiology 38:133–142.

Koelsch S, Schulze K, Sammler D, Fritz T, Muller K, Gruber O (2009): Functional architecture of verbal and tonal working memory: an FMRI study. Human Brain Mapping 30:859–873.

Krauss GL, Fisher R, Plate C, Hart J, Uematsu S, Gordon B, Lesser RP (1996): Cognitive effects of resecting basal temporal language areas. Epilepsia 37:476–483.

Langers DRM, van Dijk P (2012): Mapping the tonotopic organization in human auditory cortex with minimally salient acoustic stimulation. Cerebral Cortex 22:2024–2038.

Luders H, Lesser RP, Hahn J, Dinner DS, Morris HH, Wyllie E, Godoy J (1991): Basal temporal language area. Brain 114:743–754.

Lund JS, Yoshioka T, Levitt JB (1993): Comparison of intrinsic connectivity in different areas of macaque monkey cerebral cortex. Cerebral Cortex 3:148–162.

Mani J, Diehl B, Piao Z, Schuele SS, Lapresto E, Liu P, Nair DR, Dinner DS, Lüders HO (2008): Evidence for a basal temporal visual language center: cortical stimulation producing pure alexia. Neurology 71:1621–1627.

McDowell JE, Dyckman KA, Austin BP, Clementz BA (2008): Neurophysiology and neuroanatomy of reflexive and volitional saccades: evidence from studies of humans. Brain Cogn 68:255–270.

Melcher D, Colby CL. Trans-saccadic perception (2008): Trends in Cognitive Sciences 12(12):466–473.

Mesulam, MM, Nobre AC, Kim YH, Parrish TB (2001): Heterogeneity of Cingulate Contributions to Spatial Attention. Neuroimage 13: 1065–1072.

Moeller S, Yacoub E, Olman CA, Auerbach E, Strupp J, Harel N, Kâmil U (2011): Multiband Multislice GE-EPI at 7 Tesla, With 16-Fold Acceleration Using Partial Parallel Imaging With Application to High Spatial and Temporal Whole-Brain FMRI. Magn. Reson Med 5:1144–1153.

Moerel M, De Martino F, Formisano E (2012): Processing of natural sounds in human auditory cortex: tonotopy, spectral tuning, and relation to voice sensitivity. The Journal of neuroscience 32(41): 14205–16.

Muller RA, Kleinhans N, Courchesne E (2001): Broca’s area and the discrimination of frequency transitions: a functional MRI study. Brain Lang 76:70–76

Müri RM, Nyffeler T (2008): Neurophysiology and neuroanatomy of reflexive and volitional saccades as revealed by lesion studies with neurological patients and transcranial magnetic stimulation (TMS). Brain Cogn 68:284–292.

O’Reilly JX, Schüffelgen U, Cuell SF, Behrens TEJ, Mars RB, Rushworth MFS (2013): Dissociable effects of surprise and model update in parietal and anterior cingulate cortex. PNAS 110(38): 3660–9.

Peelen MV, Kastner S (2014): Attention in the real world: toward understanding its neural basis. Trends in cognitive sciences 18(5): 242–250.

Perrone-Bertolotti M, Kujala J, Vidal JR, Hamame CM, Ossandon T, Bertrand O, … Lachaux J-P (2012): How silent is silent reading? Intracerebral evidence for top-down activation of temporal voice areas during reading. The Journal of neuroscience 32(49):17554–62.

Pierrot-Deseilligny C, Milea D, Müri RM (2004): Eye movement control by the cerebral cortex. Curr. Opin. Neurol. 17:17–25.

Plakke B, Romanski LM (2014): Auditory connections and functions of prefrontal cortex. Frontiers in Neuroscience 8:1–13.

Platel H, Price C, Baron JC, Wise R, Lambert J, Frackowiak RS, Lechevalier B, Eustache F (1997): The structural components of music perception. A functional anatomical study. Brain 120:229–243.

Posner MI (1980): Orienting of attention. Quarterly Journal of Experimental Psychology 32:3–25.

Potter MC, Kroll JF, Yachzel B, Carpenter E, Sherman J (1986): Pictures in sentences: understanding without words. Journal of Experimental Psychology: General 115:281–294.

Purcell BA, Weigand PK, Schall JD (2012): Supplementary eye field during visual search: salience, cognitive control, and performance monitoring. The Journal of neuroscience 32(30):10273–85.

Rämä P, Poremba A, Sala JB, Yee L, Malloy M, Mishkin M, Courtney SM (2004): Dissociable functional cortical topographies for working memory maintenance of voice identity and location. Cerebral Cortex 14:768–780.

Romanski LM, Tian B, Fritz J, Mishkin M, Goldman-Rakic PS, Rauschecker JP (1999): Dual streams of auditory afferents target multiple domains in the primate prefrontal cortex. Nature Neuroscience 2:1131–1136.

Rubens AB, Kertesz A (1983): The localization of lesions in transcortical aphasias. In: Kertesz A. Localization in Neuropsychology. 1^st^ ed. Academic Press. p 245–268.

Saygin AP, Sereno MI (2008): Retinotopy and attention in human occipital, temporal, parietal, and frontal cortex. Cerebral cortex 18:2158–68.

Schiller PH, Chou IH (1998): The effects of frontal eye field and dorsomedial frontal cortex lesions on visually guided eye movements. Nature Neurosci 1:248–253.

Schira MMI, Tyler CW, Breakspear M, Spehar B (2009): The foveal confluence in human visual cortex. Journal of Neuroscience 29:9050–9058.

Schluppeck D, Glimcher P, Heeger DJ (2005): Topographic organization for delayed saccades in human posterior parietal cortex. Journal of Neurophysiology 94:1372–1384.

Schmahmann J, Pandya D (2009): Fiber Pathways of the Brain. Oxford(UK): Oxford University Press.

Sereno MI, Allman JM (1991): Cortical visual areas in mammals. In Leventhal AG. 1^st^ ed. The Neural Basis of Visual Function. London (UK). Macmillan. p 160–172.

Sereno MI, Dale AM, Reppas JB, Kwong KK, Belliveau JW, Brady TJ, Rosen BR, Tootell RB (1995): Borders of multiple visual areas in humans revealed by functional magnetic resonance imaging. Science 268:889–893

Sereno MI, Lutti A, Weiskopf N, Dick F (2013): Mapping the human cortical surface by combining quantitative T1 with retinotopy. Cerebral cortex 23:2261–2268.

Sereno MI, Pitzalis S, Martinez A (2001): Mapping of contralateral space in retinotopic coordinates by a parietal cortical area in humans. Science 294:1350–1354.

Sereno MI, Saygin AP, Hagler DJ Jr (2003): Retinotopy in parietal and temporal cortex. Neuroimage 19:S1523.

Sharp DJ, Scott SK, Wise RJS (2004): Retrieving meaning after temporal lobe infarction: the role of the basal language area. Annals of Neurology 56:836–846.

Silver MA, Kastner S (2009): Topographic maps in human frontal and parietal cortex. Trends in cognitive sciences 13:488–495.

Silver MA, Ress D, Heeger DJ (2005): Topographic maps of visual spatial attention in human parietal cortex. Journal of Neurophysiology 94:1358–1371.

Simmons W, Barsalou L (2003): The similarity-in-topography principle: reconciling theories of conceptual deficits. Cogn. Neuropsychol 20:451–486

Sperry RW (1943): Effect of 180 degree rotation of the retinal field on visuomotor coordination. Journal of experimental zoology 92:263–279.

Swisher JD, Halko MA, Merabet LB, McMains SA, Somers DC (2007): Visual topography of human intraparietal sulcus. Journal of Neuroscience 27:5326–5337.

Talavage TM, Ledden PJ, Benson RR, Rosen BR, Melcher JR (2000): Frequency-dependent responses exhibited by multiple regions in human auditory cortex. Hearing Research 150: 225–244.

Talavage TM, Sereno MI, Melcher JR, Ledden PJ, Rosen BR, Dale AM (2004): Tonotopic organization in human auditory cortex revealed by progressions of frequency sensitivity. Journal of Neurophysiology 91:1282–1296.

Thivierge J-P, Marcus GF (2007): The topographic brain: from neural connectivity to cognition. Trends in neurosciences 30:251–259.

Wandell BA, Dumoulin SO, Brewer AA (2007): Visual field maps in human cortex. Neuron 56:366–383.

Zhao M (2012): Eye movements and attention: The role of pre-saccadic shifts of attention in perception, memory and the control of saccades. Vision Research 74:40–60.

Zeharia N, Hertz U, Flash T, Amedi A (2012): Negative blood oxygenation level dependent homunculus and somatotopic information in primary motor cortex and supplementary motor area. PNAS 109:18565–18570.

